# Classification of daily crop phenology in PhenoCams using deep learning and hidden markov models

**DOI:** 10.1101/2021.10.20.465168

**Authors:** Shawn D Taylor, Dawn M Browning

**Affiliations:** US Department of Agriculture, Agricultural Research Service, Jornada Experimental Range, New Mexico State University, Las Cruces, New Mexico, 88003, USA; Oak Ridge Institute for Science and Education (ORISE), Oak Ridge, Tennessee, 37830, USA

**Keywords:** crop phenology, PhenoCam, VGG16, hidden markov model, time series, LTAR

## Abstract

Near surface cameras, such as those in the PhenoCam network, are a common source of ground truth data in modelling and remote sensing studies. Despite having locations across numerous agricultural sites, few studies have used near surface cameras to track the unique phenology of croplands. Due to management activities, crops do not have a natural vegetation cycle which many phenological extraction methods are based on. For example, a field may experience abrupt changes due to harvesting and tillage throughout the year. A single camera can also record several different plants due to crop rotations, fallow fields, and cover crops. Current methods to estimate phenology metrics from image time series compress all image information into a relative greenness metric, which discards a large amount of contextual information. This can include the type of crop present, whether snow or water is present on the field, the crop phenology, or whether a field lacking green plants consists of bare soil, fully senesced plants, or plant residue. Here we developed a modelling workflow to create a daily time series of crop type and phenology, while also accounting for other factors such as obstructed images and snow covered fields. We used a mainstream deep learning image classification model, VGG16. Deep learning classification models do not have a temporal component, so to account for temporal correlation among images our workflow incorporates a hidden markov model in the post-processing. The initial image classification model had out of sample F1 scores of 0.83-0.85, which improved to 0.86-0.91 after all post-processing steps. The resulting time series show the progression of crops from emergence to harvest, and can serve as a daily, local scale dataset of field states and phenological stages for agricultural research.

## 1. Introduction

The timing of planting, emergence, maturity, and harvest of crops affects the yield and long-term sustainability of croplands, thus tracking crop phenology has numerous interested parties from local to national levels. Remotely sensed data is an important data source for tracking crop phenology, and other attributes, from the field to global scales [1]. For example crop type, crop succession, and off-season management strategies can all be monitored using remote sensing data to inform large scale management [2]. Estimating crop phenology from remotely sensed data is difficult since crops do not follow the same patterns as natural vegetation. Harvest may happen when crops are still green, and multiple harvests in a single year will result in several “peaks” in greenness. Crop rotations result in different crop types year to year, which affects the relative greenness and derived phenology metrics [3]. A primary limitation for improving satellite remote sensing based crop phenology models is a lack of widespread ground truth data [4]. Near-surface cameras used in the PhenoCam network offer a novel solution to this need for local-scale crop management information. Images from near-surface cameras can document the date of emergence, maturity, harvest, and tillage at the field scale with a daily temporal resolution [5,6].

Automatically extracting cropland attributes and phenological information from PhenoCam imagery is difficult. The primary method to estimate phenological metrics with PhenoCam data uses the direction and amplitude of a greenness metric (the green chromatic coordinate, Gcc) of regions of interest in the camera field of view [7,8]. These metrics are well correlated with crop emergence and maturity, but cannot be used to directly identify other attributes such as crop type, flowering, or the presence of crop residue [9]. Deep learning models provide a straightforward method for identifying information in images [10], and can potentially identify phenological states directly as opposed to inferring them from relative greenness in the images. Studies have successfully used deep learning image classification models to identify and count animals [11,12], classify animal movement [13], and identify the phenological stage [14,15], species [16], or stressors [17] of individual plants. Deep learning has been used previously with PhenoCams to identify images with snow cover with up to 98% accuracy [18] and also with high accuracy to classify forest phenology [19,20].

In agriculture, deep learning classification of near surface images, either from fixed or handheld cameras, has primarily been used for weed and crop disease detection [21]. Few studies have used deep learning for the classification of crop and field attributes (e.g. [22–24]) and to our knowledge no study has applied deep learning methods at cropland sites in the PhenoCam image archive [25]. In addition to producing widespread ground truth data for large scale models, automated classification of crop phenology would be beneficial in local-scale experiments of cropping practices. Recent studies have shown phenology from PhenoCams enables high-throughput field phenotyping and tracking a variety of crop characteristics [6,26].

Here we use images from 55 agricultural cameras in the PhenoCam network to build a classification model for identifying cropland phenological states. The database includes images with a variety of real-world conditions across several crop types (Figure 1). We used a generalized classification scheme with 21 classes across 3 mutually exclusive categories, ranging from emergence to post-harvest residue. We also included classes for crop type and factors such as flooded or snow covered fields. Deep learning models designed for image classification do not have a temporal component, so we use a hidden markov model in the classification post-processing to account for the temporal correlation of daily camera time series. Results show the feasibility of a daily, local scale dataset of field states and phenological stages for agricultural research.

**Figure 1.**
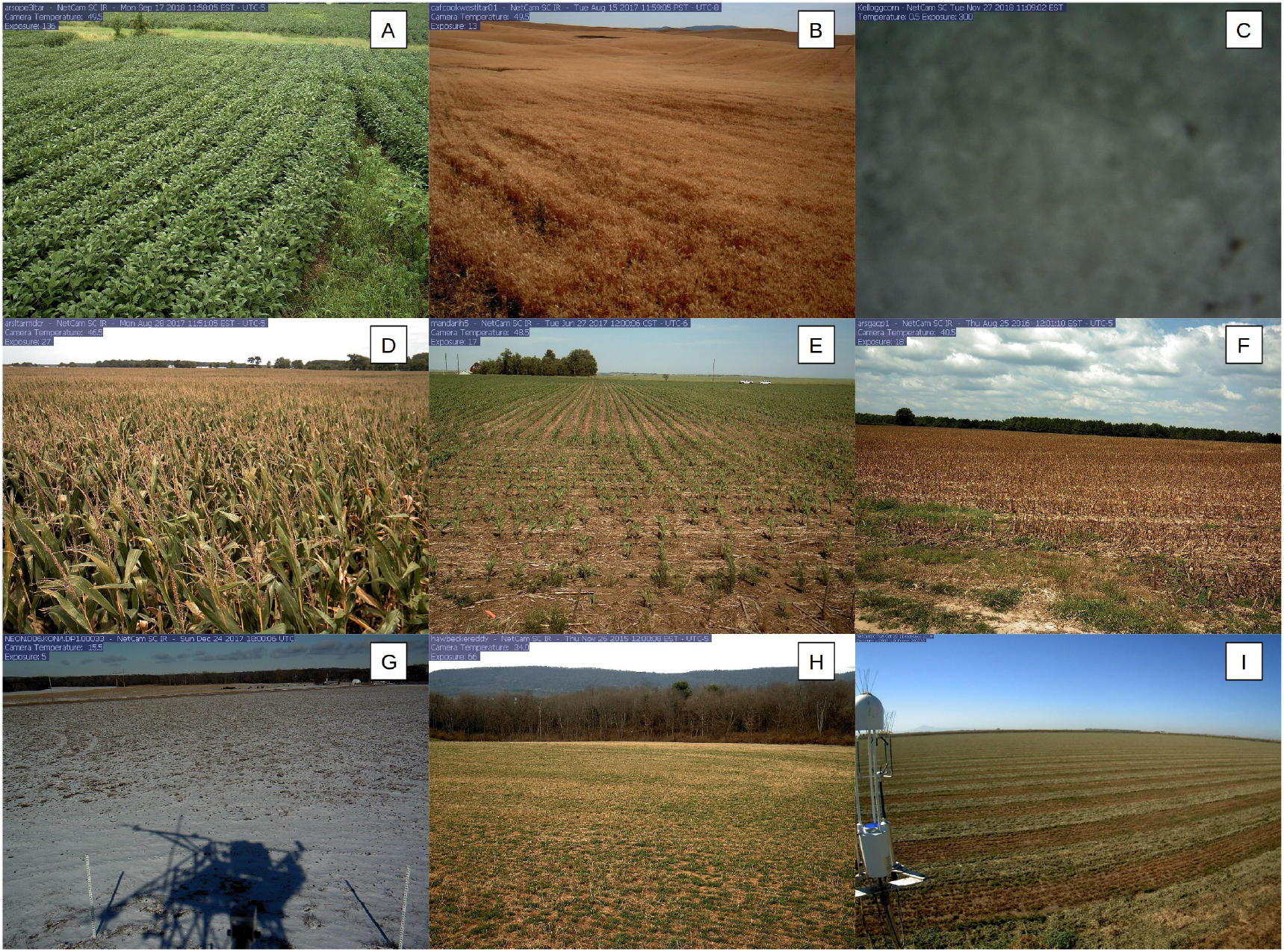
Example images from the PhenoCam network used in this study. The image sources and annotations are as follows: (A) is from arsope3ltar on Sep. 17, 2018 with soybeans in the growth stage, (B) is from cafcookwestltar01 on Aug. 15, 2017 with wheat which is fully senesced, (C) is kelloggcorn on Nov. 27, 2018 and is blurry for all categories, (D) is arsltarmdcr on Aug. 28, 2017 with corn in the flowers stage, (E) is from mandanh5 on June 27, 2017 and has corn in the growth stage with a dominant cover of residue, (F) is from arsgacp1 on Aug. 25, 2016 with dominant cover of plant residue and no crop present, (G) is from NEON.D06.KONA.DP1.00033 on Dec. 24, 2017 with dominant cover of snow and no crop present, (H) is from hawbeckereddy on Nov. 26, 2015 with an unknown crop type in the growth state, and (I) is from bouldinalfalfa on Oct. 22, 2019 with a dominant cover of residue and no crop present. Images A, B, D, and H all have vegetation as the dominant cover class.

## 2. Methods

### 2.1. Data

We used PhenoCam images from agricultural sites to train an image classification model (Figure S1, Table S1). To obtain a representative sample of images across all potential crops and crop stages we used seasonal transition dates provided by the PhenoCam Network. Based on the transition date direction (either rising or falling) and threshold (10%, 25%, and 50%) we partitioned each calendar year into distinct periods of senesced, growth, peak, and senescing [7]. We chose 50 random days from each site, year, and period, for a total of 8,270 images. We annotated each image by hand using the imageant software into the 21 classes described below [27].

Initial image classifications were organized into 21 classes across three categories of Dominant Cover, Crop Type, and Crop Status (Table 1). The categories are each mutually exclusive such that any single image can be independently classified into a single class within each category. This allows finer grained classification given an array of Crop Types, and more flexibility in classifying crop phenological stages. For example, it would be informative for remote sensing models to know the exact date of crop emergence, but also that on the specified date and for several days to weeks after the feld is still predominantly bare soil. The first category, Dominant Cover, is the predominant class within the field of view. The Crop Type category represents the four predominant crops in the dataset (corn, wheat/barley, soybean, and alfalfa). Wheat and barley are combined into a single category as they are difficult to discern in images. The unknown Crop Type class is used during emergence when an exact identification is impossible. The other Crop Type class represents all other crops, including fallow fields, besides the four predominant ones. The stages of the Crop Status category are loosely based on BBCH descriptions [28], but generalized to be applicable across a variety of crops and what is discernible. The Flowers stage is used solely for identifying reproductive structures for corn, wheat, or barely. No reproductive structures were discernible in images with Crop Types of Soybean, Alfalfa, or Other, thus images in those three Crop Type classes have no Crop Status annotations of Flowers.

**Table 1:** Class descriptions used in the classification model.

| Category | Class | Used in final product | Description |
| --- | --- | --- | --- |
| Dominant Cover | Blurry | No | Image blurry, out of focus, or otherwise obscured. |
|  | Vegetation | Yes | Live or senesced vegetation |
|  | Residue | Yes | Post-harvest plant residue |
|  | Bare soil | Yes |  |
|  | Snow | Yes |  |
|  | Water | Yes |  |
| Crop Type | Blurry | No | Image blurry, out of focus, or otherwise obscured. |
|  | Unknown Plant | Yes | Plants are present but cannot be confidently identified |
|  | Corn | Yes |  |
|  | Wheat/Barley | Yes |  |
|  | Soybean | Yes |  |
|  | Alfalfa | Yes |  |
|  | Other | Yes | Any other crop or a fallow field. |
|  | No crop | Yes | No crop present (eg. a plowed field or completely snow covered) |
| Crop Status | Blurry | No | Image blurry, out of focus, or otherwise obscured. |
|  | Emergence | Yes | First shoots and/or leaves are visible. |
|  | Growth Stage | Yes | Plants have several distinct leaves and/or tillers visible, but no visible tassels, flowers, or fruit. |
|  | Tassels/Flowering | Yes | Plants have distinct tassels, flowers, or fruit. |
|  | Senescing/browning | Yes | 10% or more of visible plants are brown/browning. |
|  | Fully senesced | Yes | 90% or more of visible plants are fully senesced. |
|  | No crop | Yes | No crop present (eg. a plowed field or completely snow covered) |

After annotation we excluded some images based on low prevalence of some category combinations. For example, only 8 images had the combined combination of Soil, Unknown Plant, and Senescing for the Dominant Cover, Crop Type, and Crop status categories, respectively. When a unique combination of the three categories had less than 40 total images, all images representing that combination were excluded from the model fitting. This resulted in 255 annotated images, from the original 8,270, being excluded. A total of 8,015 annotated images were available for model fitting.

We used mid-day images in the annotation stage and leveraged more of the PhenoCam archive to increase sample size for model fitting. The 8,015 images that we annotated represent the midday image for a single date, though phenocams record images up to every 30 minutes. For each annotated image date we also downloaded all images between 0900 and 1500 local time, resulting in an additional 83,469 images. We applied the annotation of the midday image to all images of that date, resulting in 91,484 total images used in the model fitting. This allowed us to increase training image data by a factor of 10 with minimal effort, and include more variation in lighting conditions. While it’s possible some of these non-midday images were annotated incorrectly (e.g., a blurry camera becoming cleared, or a field being plowed after midday) these are likely minimal and did not have a large effect on model accuracy [12].

### 2.2. Image Classification Model

We used the VGG16 model in the Keras python package to classify images into the 21 classes (Table 1)[29,30]. We evaluated several other models preconfigured in the Keras package and found VGG16 to be the best performing (see Supplement). The model allows us to specify the hierarchical structure of the three categories, such that the predicted class probabilities for any image sum to one within each category. We held out 20% of the images as a validation set. The validation set included all images from three cameras: arsmorris2, mandani2, and cafboydnorthltar01 totaling 10,172 images. It also included 8,124 randomly selected images from the remaining locations to obtain the full 20%. This resulted in a validation sample size of 18,296 and a training sample size of 73,188. The VGG16 classification model was trained fully, as opposed to using transfer learning [12]. We experimented with transfer learning, where a pre-trained model is fine-tuned using our own data, but found that training the model fully had better results.

We resampled the 73,188 training images to 100,000 using weights proportional to the unique combinations among the three categories. For example, there were 4,407 images annotated as Vegetation, Corn, and Flowers for the three categories, but only 1,074 images annotated with Vegetation, Wheat, and Flowers. The images in the former class were given a lower weight in the resampling to reach 100,000 total training images. This allows for even sample sizes among classes and protects against the model being biased toward common classes. During model fitting the images are shuffled and transformed using random shifts and rotations such that the exact same image is never seen twice, which protects against over-fitting. We trained the model with an image resolution of 224×224 pixels using the Adams optimizer with a learning rate of 0.01 for 15 epochs, and an additional 5 epochs with a learning rate of 0.001.

### 2.3. Post-processing

After fitting, the VGG16 model was used to classify approximately fifty-five thousand midday images from all agricultural PhenoCam sites, totalling approximately 170 site-years [31]. These classified images were then put through a post-processing routine to produce a final classification for each day (Table 2). First, for all dates marked as Snow in the Dominant Cover category, the Crop Type and Crop Status predictions were removed and gap-filled using linear interpolation from surrounding non-snow dates as long as the gap was 60 days or less. The reasoning behind this is during the constant snow cover of winter the crop, if any, likely remains unchanged. Next, any image marked as blurry was removed and the associated image date marked as missing across all three categories (Dominant Cover, Crop Type, Crop Status). Gaps of missing dates, up to 3 days, in the time series for each site were filled with a linear interpolation of the two bounding date probabilities for each of the remaining 18 classes. Probabilities across all dates were normalized across the remaining classes to account for removing the “Blurry” class. After this initial filtering the classification time series from the approximately fifty-five thousand images were input into an hidden markov, and several additional post-processing steps, to produce the final time series.

**Table 2:** The post-processing steps performed on the VGG16 model predictions.

|  |
| --- |
| 1. Predict probabilities for each of the 55k daily images. |
| 2. For snow days remove Crop Type and Crop Status and gap-fill up to 60 days. |
| 3. For blurry images remove all predictions for that date and gap fill up to 3 days. |
| 4. Apply HMM to Dominant Cover and Crop Status categories. |
| 5. Identify each unique crop sequence. A crop sequence is all dates between two non-consecutive “no crop” dates of the Crop Status category. |
| 6. For each crop sequence identify the Crop Type using the highest cumulative probability excluding the unknown class. |
| 7. Mark crop sequences as unknown for sequences 60 days or less, where the most common Crop State was emergence. |
| 8. Mark crop sequences as unknown when the highest cumulative probability within the sequence was No Crop. |

Out-of-the box image classification models such as VGG16 have no temporal component. Every image is treated as an observation independent of temporally adjacent images. Thus mis-classification of images can lead to noisy time series and improbable transitions between classes. To correct for this we used a hidden markov model (HMM) to reduce the day-to-day variation and remove improbable transitions [32–34]. HMM’s are state-space models which combine a latent “true” state of a process with an observation model. The latent state evolves dynamically where every timestep is a discrete state which depends only on the state of the previous timestep, and where the probability of moving from one state to another is decided by a transition matrix. The observation model is a timeseries of the same length where every observed state depends only on the latent state of the same timestep. Given a latent state, all observed states have a non-zero probability of being observed.

We used two HMMs, one for Dominant Cover and another for Crop Status, each with a daily timestep. Only sequences with at least 60 continuous days, after the gap-filling from Snow and Blurry dates described above, were processed with the HMM. For the observation model within each HMM we used the direct output from the classification step, which for each image date consisted of probabilities of the image belonging to each class in the respective category. The transition matrix describing the probabilities of the hidden state changing states from one day to the next was created manually for each HMM (Tables S2-3). The probability of the hidden state staying the same between two dates was set as the highest (0.90 - 0.95) in all cases. In this way the hidden state only changes when there is strong evidence in the observation model, represented by continuous and high observed probabilities of a new state. Within the Dominant Cover category, transitions to other states besides the current one were set to equal values with one exception. Transition from Soil to Residue was set to 0 probability since residue can only be present after a crop has been harvested. Within Crop Status transitions between states were constrained to be biologically possible. Improbable transitions (e.g., from Emergence to Senescing in a single day) had probabilities of 0. Reproductive structures were not visible on all crops in images, so transitioning from the Growth stage directly to Senescing was allowed. Transitioning from either Senescing or Senesced to Growth was also allowed since this represents dormancy exit in overwintering crops such as winter wheat. Given observation probabilities and the transition matrices the most likely hidden state was predicted using the viterbi algorithm to produce the final time series across the Dominant Cover and Crop Status categories.

For the Crop Type category the HMM methodology is less useful, since no day-to-day transitions between Crop Types are expected. Here we used the HMM output of Crop Status to identify each unique crop sequence, defined as all dates between any two “No Crop” classifications. For each unique crop sequence, we identified the associated Crop Type with the highest total probability within that sequence, and marked the entire sequence as that Crop Type. In this step we considered all Crop Type classes except “Unknown Plant”, which is used during the emergence stage when a plant is present but the exact type is unclear. This allows information later in the crop cycle, when Crop Type is more easily classified, to be propagated back to the emergence stage. Finally, we marked the final Crop Type as Unknown for some crop sequences in two instances: 1) when the length of a crop sequence is less than 60 days and the sequence was predominantly in the emergence stage, and 2) when the “No Cop” class was selected as the final Crop Type with the highest probability within a sequence. These two scenarios tended to occur when volunteer plants are growing sparsely on an otherwise bare field.

### 2.4. Evaluation

We calculated three metrics to evaluate the performance of the image classifier: precision, recall, and the F1 score. All three metrics are based on predictions being classified into four categories of true positive (*TP*), false positive (*FP*), true negative (*TN*), and false negative (*FN*). Precision is the probability that an image is actually class *i*, given that the model classified it as class *i*. Recall is the probability that an image will be classified into class *i*, given that the image is actually class *i*. The F1 score is the harmonic mean of precision and recall. All three metrics have the range 0-1, where 1 is a perfect classification.

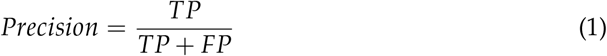

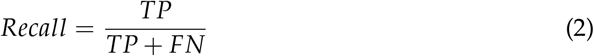

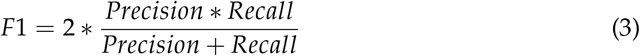

where *TP*, *FP*, *FN* are the number of true positive, false positive, and false negative classifications, respectively. Using only the mid-day images we calculated all three metrics for the 21 original classes to evaluate the performance of the VGG16 model. For this step the predicted class for each category on a single date was the one with the maximum probability. We re-calculated all three metrics again after all post-processing steps. The second round of metrics does not include two classes removed during post-processing: the Blurry class across all categories and the Unknown Plant class in the Crop Type category.

Software packages used throughout the analysis include Keras v2.4.3 [30], TensorFlow v2.2.0 [35], Pandas v1.0.4 [36], NumPy v.1.20.2 [37], and Pomegranate v0.14.5 [38] in the Python programming language v3.7 [39]. In the R language v4.1 [40] we used the zoo v1.8.0 [41], tidyverse v1.3.1 [42], and ggplot2 v3.3.5 [43] packages. All code for the analysis, as well as the final model predictions, are available in a Zenodo repository ([44], https://doi.org/10.5281/zenodo.5618316).

## 3. Results

The overall F1 score, a summary statistic which incorporates recall and precision, was 0.90-0.92 for the training data across the three categories of Dominant Cover, Crop Type, and Crop Status (Figure 2). The overall F1 score for validation data, which was not used in the model fitting, was 0.83-0.85 for the three categories. In the Dominant Cover category the vegetation class was the best performing overall with recall and precision of 0.97 and 0.93, respectively. Thus the classification model has a strong ability to discern when the camera field of view is or is not predominantly vegetation. When vegetation is not dominant the classifier is still moderately accurate, though there is confusion between soil and residue classes indicated by their recall scores (0.64-0.68). The precision of soil and residue was 0.61 and 0.82 for validation data, indicating that the classifier leaned toward residue.

**Figure 2.**
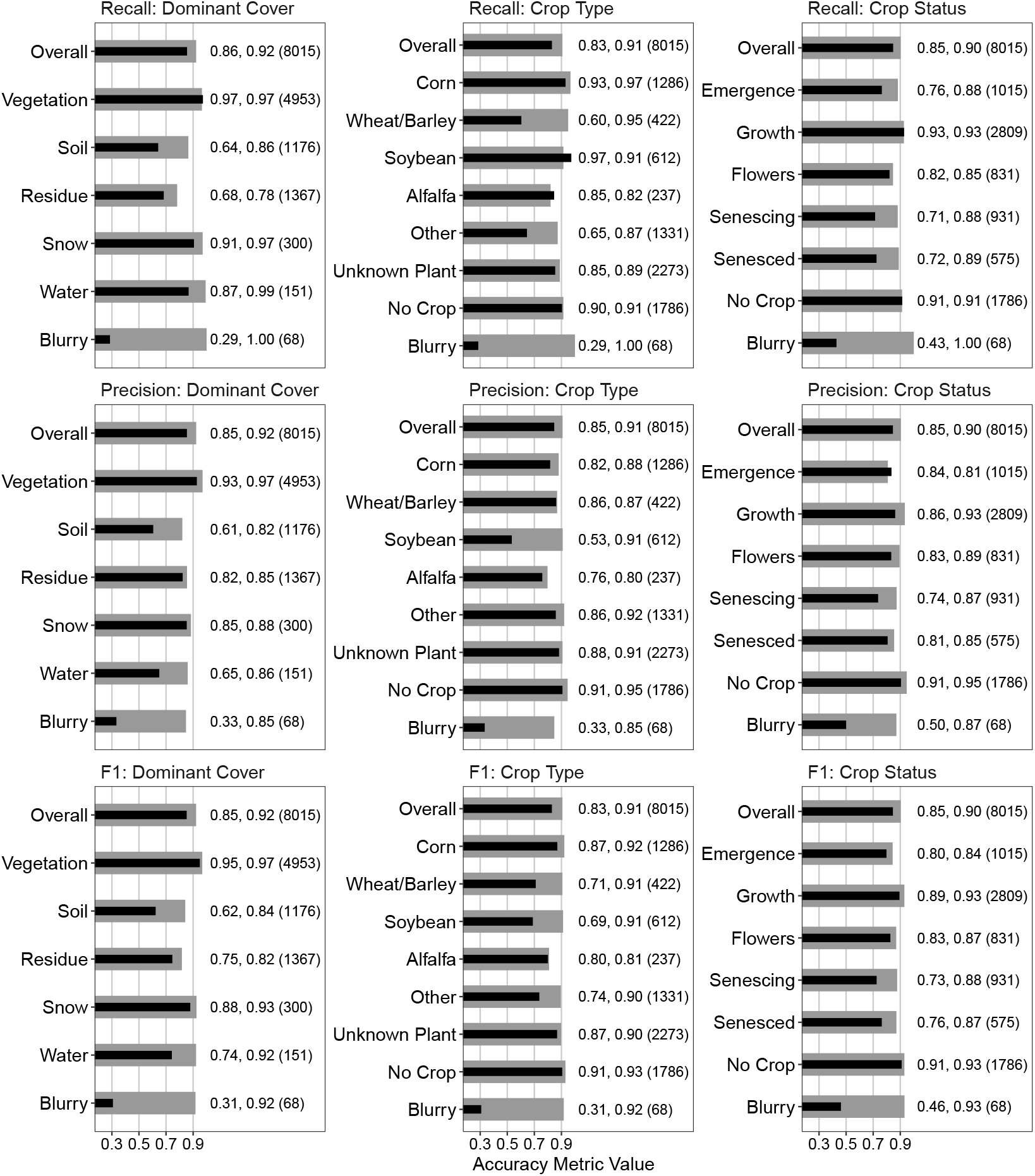
Accuracy metrics for the VGG16 image classifier. The black and grey bars represent the validation and training datasets, respectively. The text indicates the respective metric value for validation, training, and class sample size in parentheses. The training and validation sample sizes are 80% and 20% of the total sample size, respectively. Overall indicates the average metric value for the respective category, weighted by sample size. All three metrics have a range of 0-1 where 1 equals a perfect prediction.

Excluding the blurry class, the worst precision for Crop Type was soybeans, with a precision of 0.53 on the validation data. This indicates that if an image was classified as Soybean, then there is a 53% chance it is actually soybean. The recall for Soybean was high, 0.97 with the validation data, indicating that there is a high amount of false positives from non-Soybean images being classified as Soybean. Conversely the Wheat/Barley and Other classes have high precision (0.86 and 0.86, respectively), and low recall (0.60 and 0.65, respectively). This indicates a high amount of false negatives, where images of Wheat/Barley and Other Crop Types are being classified as other Crop Type classes (Figure S2,S3).

The blurry class had low recall and precision across all three categories, with values of 0.29-0.43 and 0.33-0.50 for validation data recall and precision, respectively. Combined with training data recall scores of 1.0, this indicates likely over-fitting of the blurry class in the classification model. We did not attempt to improve this further since the blurry image prevalence was extremely low. Additionally, when images were marked as blurry in the final dataset the final state was interpolated in the HMM post-processing step by accounting for the surrounding images.

Figure 3 shows the classification statistics after post-processing of the image time series, where the HMM was used for Dominant Cover and Crop Status and the Crop Type was set to the highest total probability within any single crop series. The blurry class is not shown here since it was removed in the post-processing routine. The Unknown Plant class for Crop Type is also excluded since in the post-processing the Crop Type category is assigned to the highest probability class seen in each crop sequence, thus performance metrics for the Unknown Plant class would be uninformative.

**Figure 3.**
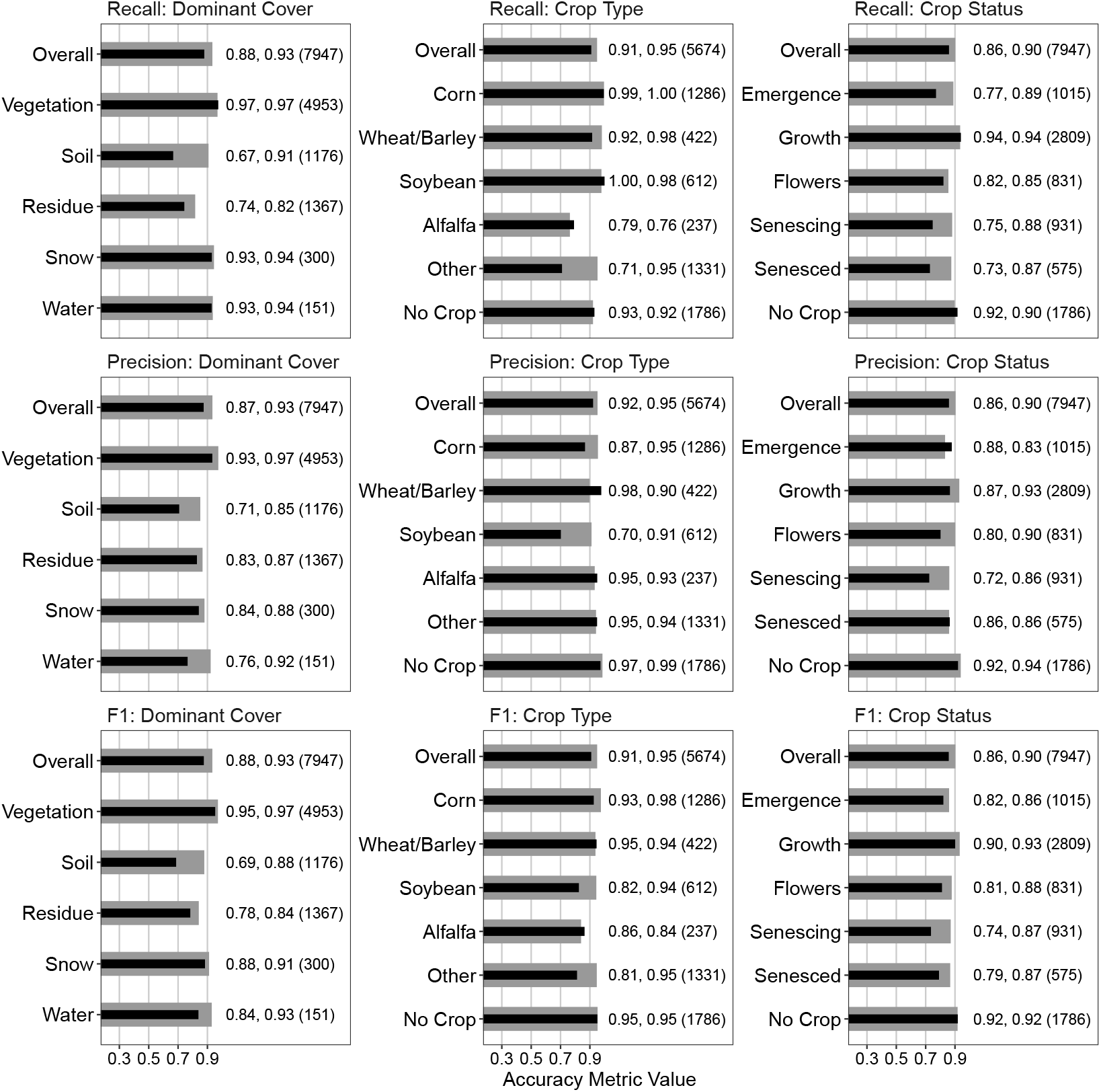
Accuracy metrics for the classifications after post-processing. The black and grey bars represent the validation and training datasets, respectively. The text indicates the respective metric value for validation, training, and class sample size in parentheses. The training and validation sample sizes are 80% and 20% of the total sample size, respectively. Overall indicates the average metric value for the respective category, weighted by sample size. All three metrics have a range of 0-1 where 1 equals a perfect prediction. Differences between this and Figure 2 is the exclusion of the blurry and unknown plant classes, with total sample sizes reflecting this.

The validation data performance metrics after the post-processing steps either improved or remained the same across all classes except four: the Snow class in Dominant Cover, the Alfalfa Crop Type, and Flowers and Senescing Crop status classes. Overall F1 scores using validation data increased from 0.85 to 0.88, 0.83 to 0.91, and 0.85 to 0.86 for Dominant Cover, Crop Type, and Crop Status categories, respectively.

Next we present four examples of the full classification and post-processing results using a single calendar year from four sites. They show the initial output of the classification model, as well as the capability of the HMM in removing high variation in the original VGG16 model prediction. The original VGG16 predictions are obtained by choosing the class with the maximum probability (MaxP) for each category and day. We compare them with insight gained from the full image time series available on the PhenoCam data portal https://phenocam.sr.unh.edu. For example at the arsmorris2 site in central Minnesota, in the months March through June of 2020, there is uncertainty in whether the Dominant Cover of the field is Residue or Soil (Figure 4A, MaxP). The HMM model resolved it to the Residue class for the three month period. From mid-June thru October there is high certainty that that vegetation is present, reflected in both the initial classifications (MaxP) and resulting HMM. The HMM model resolved the Crop Type as Corn (Figure 4B). During June to October, the Crop Status progresses naturally through the different stages, and uncertainty arises only in October when fully senesced vegetation is confused with plant residue (Figure 4C).

**Figure 4.**
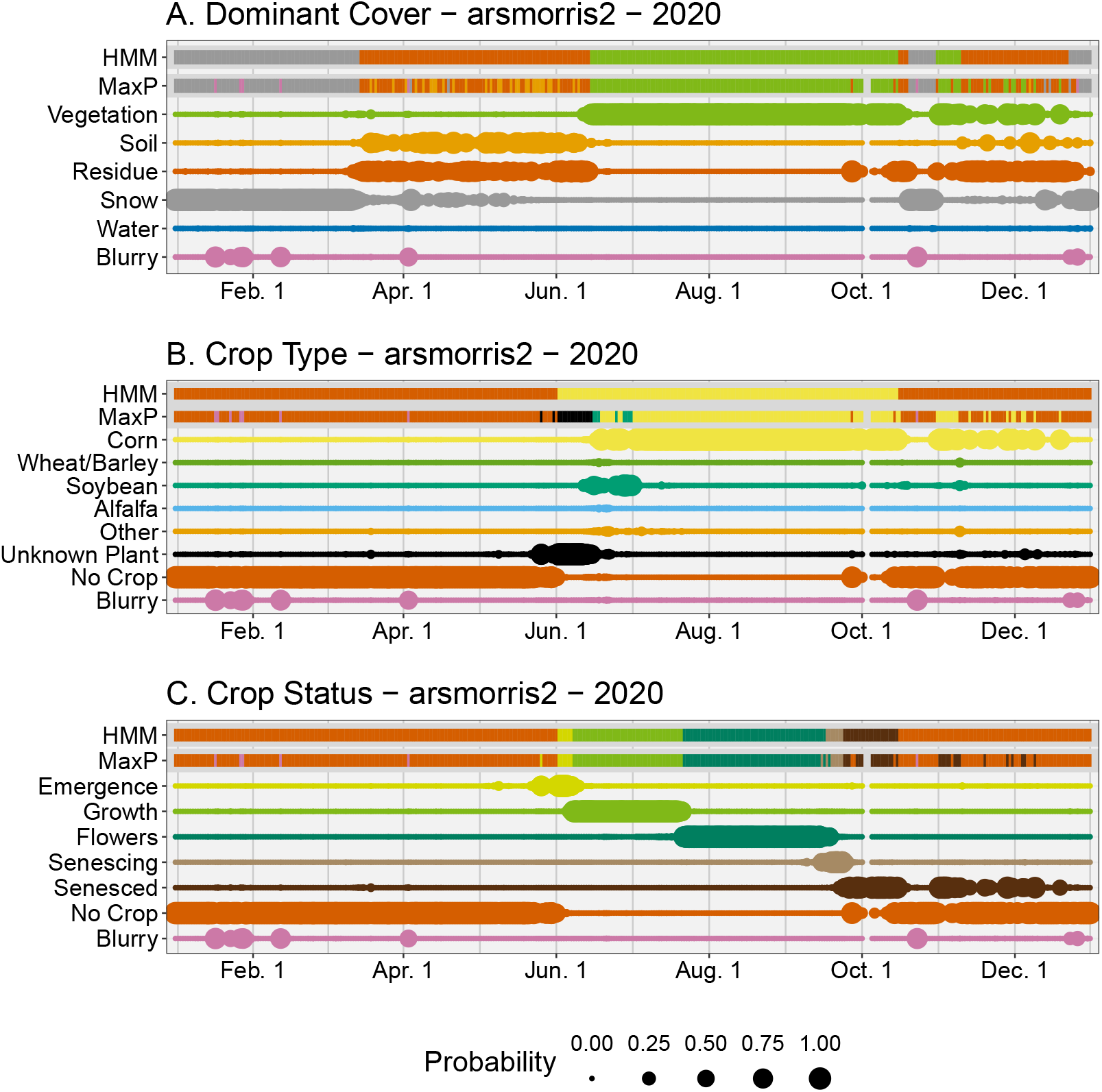
Classification results for the arsmorris2 site for the year 2020. The panels represent the results for the Dominant Cover (A), Crop Type (B), and Crop Status (C) categories. The top two rows of each panel represent the final classification for either the daily maximum probability (MaxP) or the hidden markov model (HMM). The remaining rows in each panel represent the initial model classification for the respective class, where larger sizes represent higher probability.

Cropping systems with multiple harvests per season are challenging for remote sensing models. There were multiple harvests for the bouldinalfalfa site in northern California for the year 2018. Here an alfalfa field was persistent for the entire year with several harvests (Figure 5A). During the intervals of regrowth after each harvest the Dominant Cover of the field was classified as Residue with emergence of an Unknown crop type (Figure 5B). Once the plants matured then it was identified consistently as Alfalfa, which in the post-processing was back propagated in time for each crop sequence.

**Figure 5.**
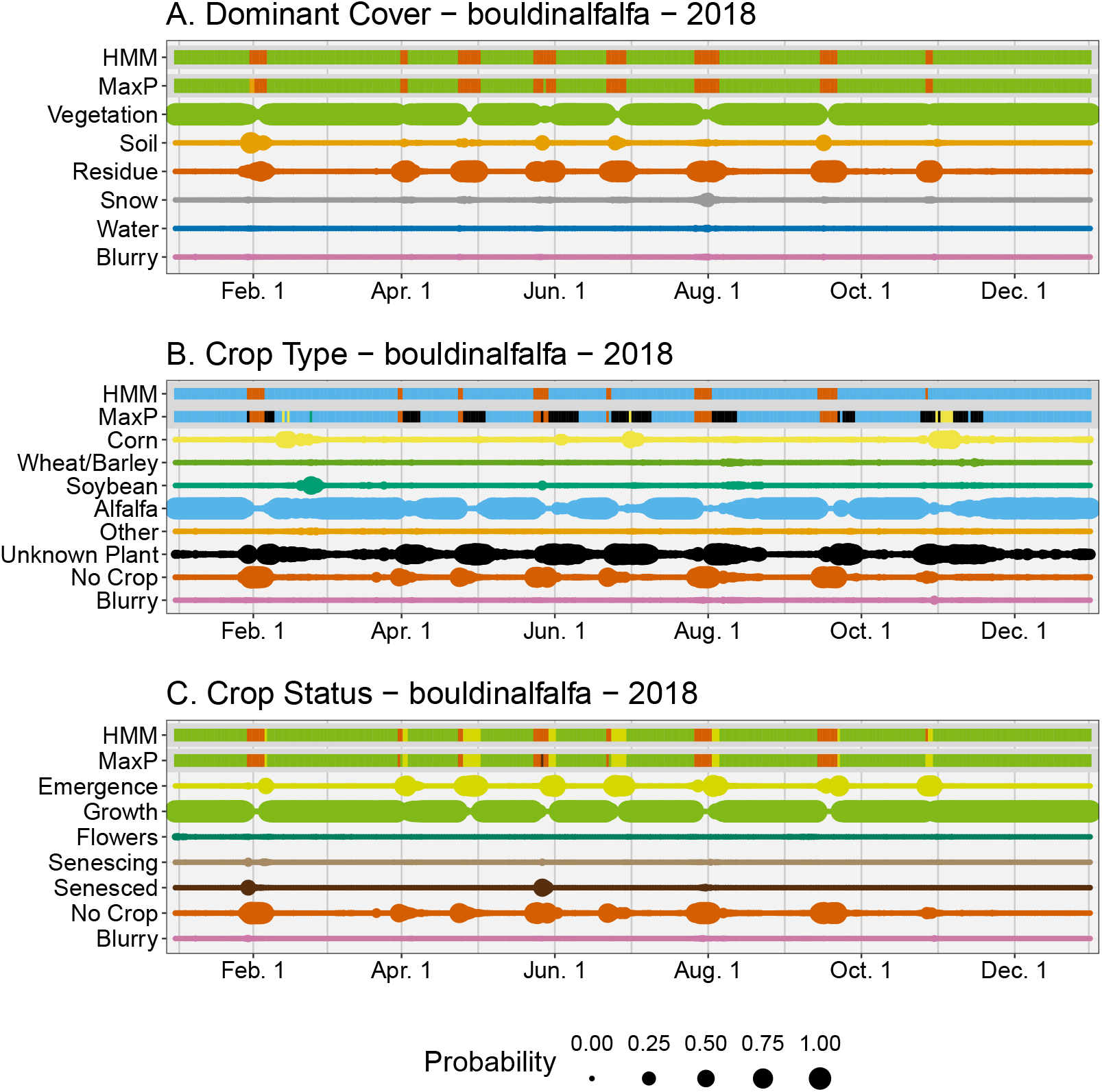
Classification results for the bouldinalfalfa site for the year 2018. The panels represent the results for the Dominant Cover (A), Crop Type (B), and Crop Status (C) categories. The top two rows of each panel represent the final classification for either the daily maximum probability (MaxP) or the hidden markov model (HMM). The remaining rows in each panel represent the initial model classification for the respective class, where larger sizes represent higher probability.

At the site cafcookeastltar01 in eastern Washington in the year 2018 there was a short residual crop of wheat in April (Figure 6A). Since the plants were not allowed to grow into the summer, due to a new crop being planted, they were not positively identified and instead marked as Unknown Plant. From manual image interpretation, we know a crop of chickpeas was planted in May which grew until harvest in early September. Throughout the summer the model initially classifies this crop as wheat, soybean, or alfalfa. The post-processing correctly chose the Other Crop Type as the final class. In October and November there is confusion in the Dominant Cover category between soil and residue, even after post-processing. From the images we can conclude there was likely no activity in the field during this time, thus confusion likely stems from a moderate amount of residue on the field combined with low light conditions of this northern (47.7° latitude) site.

**Figure 6.**
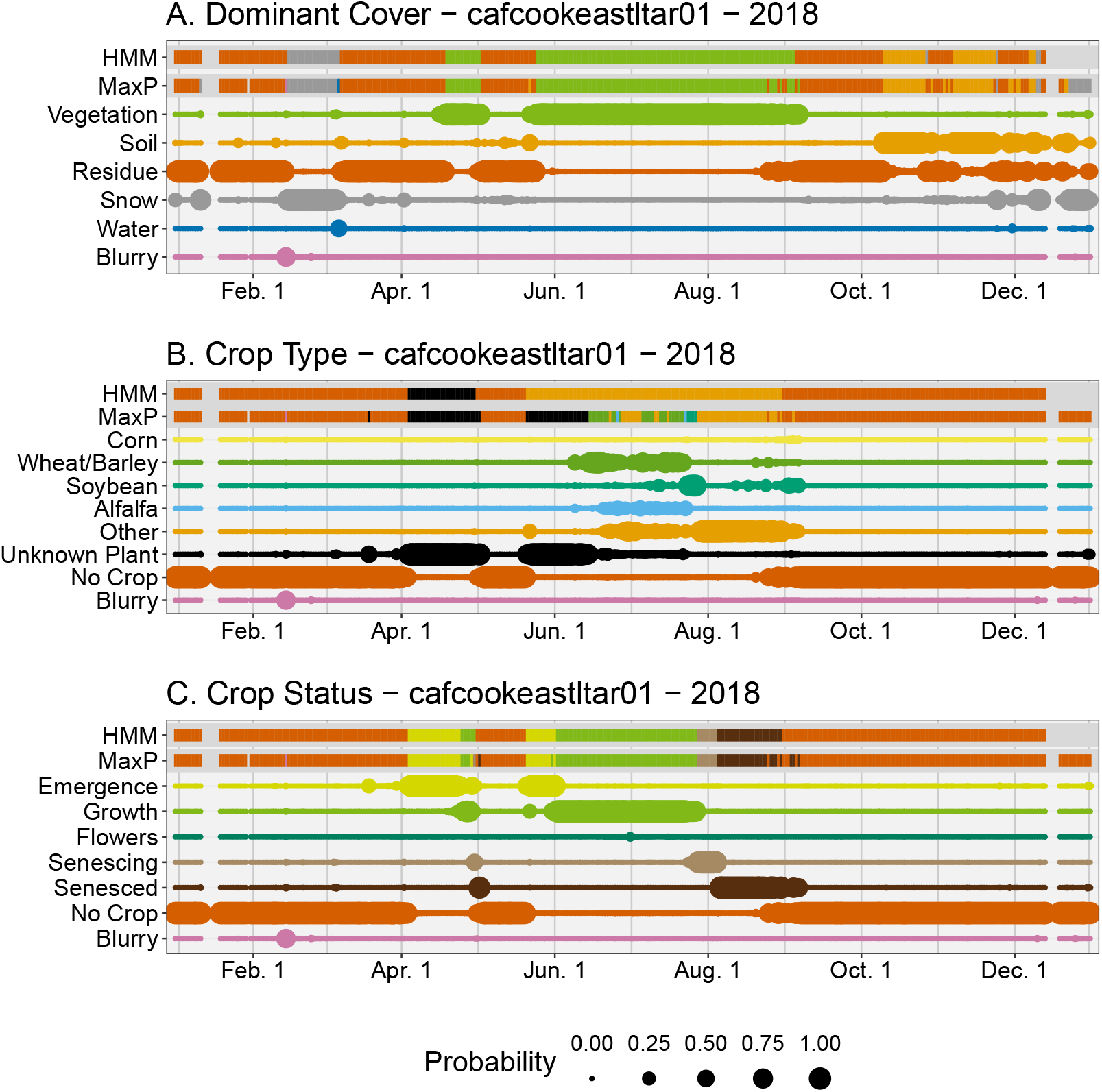
Classification results for the cafcookeastltar01 site for the year 2018. The panels represent the results for the Dominant Cover (A), Crop Type (B), and Crop Status (C) categories. The top two rows of each panel represent the final classification for either the daily maximum probability (MaxP) or the hidden markov model (HMM). The remaining rows in each panel represent the initial model classification for the respective class, where larger sizes represent higher probability.

Crops going into dormancy in the winter and resuming growth in the spring are accounted for in the post-processing routines as demonstrated by the Konza Agricultural site in the NEON network in 2017 (NEON.D06.KONA.DP1.00042, Figure 7). A winter wheat crop (Figure 7B), which was planted in the fall of 2016, resumed growth in February. The remainder of that crop life cycle proceeded normally until harvest in July (Figure 7B). The crop type here is correctly classified as Unknown Plant by the classifier from January thru March here, since the plants were relatively small at this time. The correct classification of wheat began in March when the plants were large enough to confidently identify, and this was propagated back to the initial emergence in 2016. Additionally, the primary field at this site was harvested at the end of June 2017 (as seen in the original images), though the classification model indicated it happened mid-July (Figure 7A). This was due to the foreground plants being removed in mid-July, while the primary field was harvested in June.

**Figure 7.**
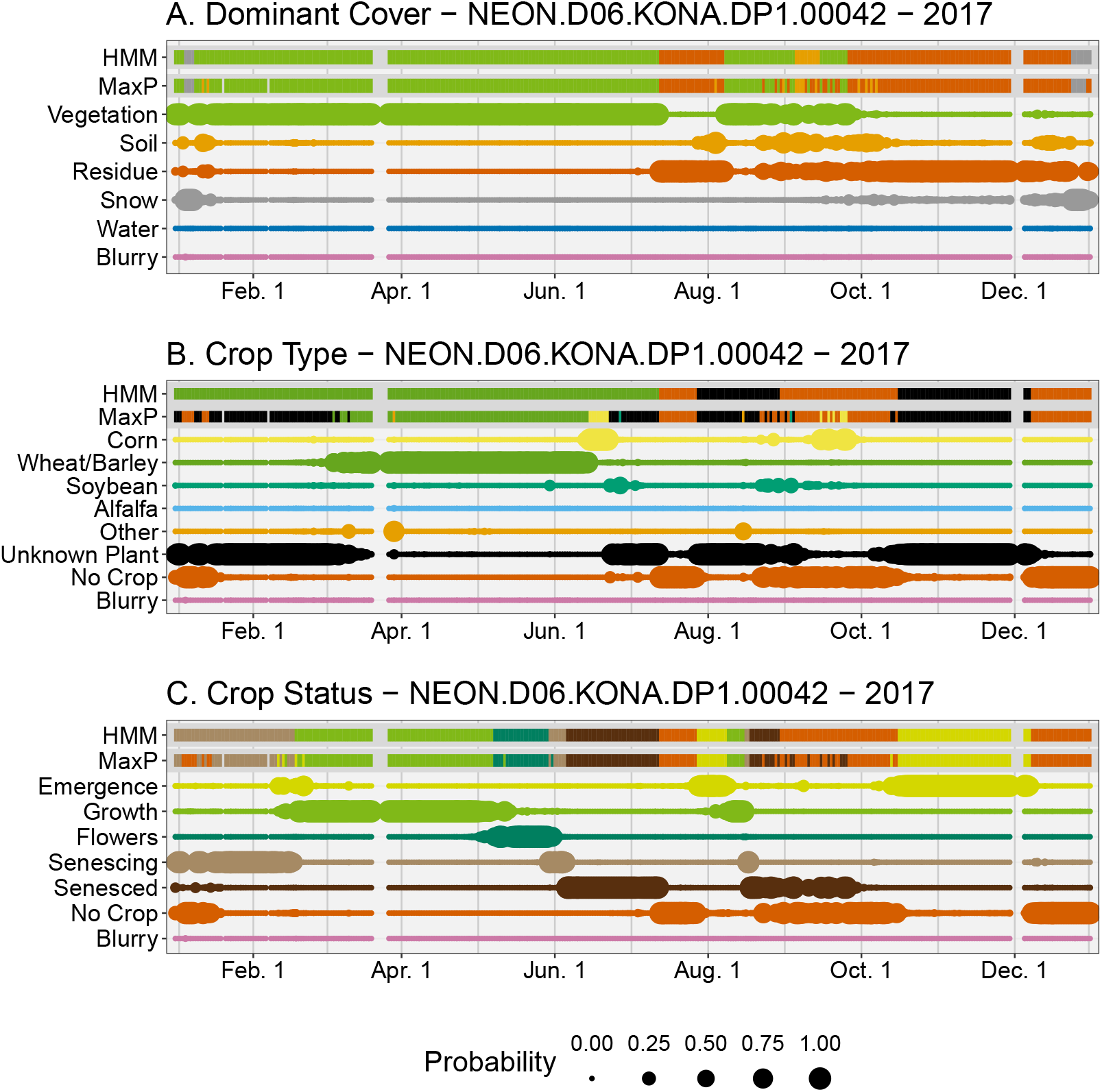
Classification results for the NEON.D06.KONA.DP1.00042 site for the year 2017. The panels represent the results for the Dominant Cover (A), Crop Type (B), and Crop Status (C) categories. The top two rows of each panel represent the final classification for either the daily maximum probability (MaxP) or the hidden markov model (HMM). The remaining rows in each panel represent the initial model classification for the respective class, where larger sizes represent higher probability.

## 4. Discussion

Daily images from the PhenoCam network contain a wealth of information beyond just vegetation greenness, and here we showed they are also a novel source of cropland phenological information. Using a deep learning-based image classification model we identified the daily field state, crop type and phenological state from PhenoCam images in agricultural fields. Since mainstream classification models do not have a temporal component we applied a hidden markov model as a post-classification smoothing method which accounts for temporal correlation. This improved classification metrics and removed improbable transitions. Improvements would be beneficial to better classify field and crop states outside the primary growing season, and to better account for crops which go through a period of dormancy.

The classification model here was developed to simultaneously identify several crop and field attributes and has a variety of potential uses. For example the United States Department of Agriculture currently monitors crop status throughout the USA using surveys [45]. An array of PhenoCams positioned in representative fields could enable a real-time crop status monitoring system using the methodologies outlined here. Remote sensing models for monitoring crop progression would benefit from the large temporal and spatial extent of PhenoCams in agricultural fields as a source of verification data [4]. The on-the-ground daily crop status data could also be used to parameterize or validate earth system models, where crop phenology is a primary source of error for crop yields [46].

Numerous studies have used *Gcc* from PhenoCams to study various biological processes [25,47]. Yet compressing images down to a single greenness index discards large amounts of information, especially in agricultural fields which are constantly managed [9]. Our approach here allows us to extract more relevant data from images, such as the crop type or the state of the field after vegetation is removed. These image-based metrics of crop type and stage from PhenoCam time series can complement *Gcc* as opposed to replacing it though. For example, *Gcc*, and other greenness metrics, can be used to derive the date of peak greenness or the rate of greenup or greendown, which reflects important plant properties not present in our image classification approach such as water stress and plant vigor [6,26,48]. Added insight from complementary metrics enrich interpretation and offer decision-makers flexibility in crop management [9].

Most studies using deep learning methods to identify cropland attributes use satellite or aerial imagery [21], though several studies have used near-surface imagery similar to the work here. Yalcin 2017 used a CNN to classify crop types and had F1 scores of 0.74-0.87. Han et al. 2021 used a CNN to classify development stages in rice with F1 scores ranging from 0.25-1.0. The high accuracy seen here and in other studies shows the capability of tracking crop and field attributes with near-surface cameras. This approach is advantageous since the cameras are not affected by cloud cover and, after initial installation, do not have significant labor costs.

We identified several areas of our approach which could be improved. Firstly, the VGG16 model used here could be replaced with either a more advanced or a customized neural network model. Though the initial accuracy of the VGG16 model was relatively high, it was originally designed for classification of common objects as opposed to croplands. It could likely be improved through model customization or fine-tuning of parameter estimation. Improving the initial image classification would improve the final results without any other adjustments to the post-processing routines.

Our approach here worked best during continuous periods where crops were present on the field. Once crops were removed, the dominant cover state could be difficult to discern due to soil, plant residue, and fully senesced plants having similar visual characteristics (Figures S4-S11). Improvements could potentially be made here by using a zoomed-in or cropped photo of the field. Since the images were compressed from their full resolution to 224×224 pixels, it is likely important details were lost. Han et al. 2021 showed that zoomed-in images, used simultaneously with full resolution images in a custom neural network model, greatly improved accuracy of rice phenology classification. Using zoomed-in images may also help with identifying the reproductive structures of crops. Though this may be limited by camera placement since even during manual annotation we could only identify reproductive structures of corn, wheat, and barley. Residue versus Soil classification may also be improved by classifying the amount of residue (e.g., the fractional cover of plant residue or soil) as opposed to using two distinct classes.

Our use of an HMM is an ideal solution to account for temporal correlation in the classified image time series. The progression of crops at a daily time step is constrained by plant biology, and these constraints are easily built into the HMM using the transition matrix. Additionally, the predicted probabilities from the classification model can be used directly in the HMM observation model, resulting in a straightforward data workflow. Since we used a basic HMM we had to create separate models for the Dominant Cover and Crop Status categories, which resulted in occasional inconsistencies. For example the Dominant Cover HMM may occasionally identify a time period as being predominantly vegetation, while the Crop Status HMM identifies the same time period as having no crop present (Figure 7). A multi-level, or layered, HMM may be able to overcome this by modelling the joint probabilities of classes across the two categories [49]. Temporal segmentation, a newer deep learning approach which is under active development, could model the joint probability of classes across the different categories in addition to having better performance than seen here [50]. A downside to temporal segmentation is that it would require fully annotated training sequences (i.e., annotations for all images in a year for numerous sites) as opposed to the random selection of training images used here.

We observed some mis-classifications when field management activities are not uniformly applied to all parts of a field in the camera field of view. In the NEON-KONA example (Figure 7) the final classification showed vegetation present due to foreground plants remaining even though the primary field was harvested. This could be improved by having a pre-processing step which identifies distinct agricultural fields within the camera field of view. Each agricultural field could then be classified independently. This would also allow the inclusion of PhenoCam sites focused on one to several experimental plots, which were excluded from this study. This step could be done automatically through image segmentation models, or manually as in the region of interest (ROI) identification in the current PhenoCam Network data workflow [7].

Instead of discarding blurry or obscured images we accounted for them directly in the modelling. This is ideal since real-time applications must account for such images without human intervention. The Blurry class across all three categories had high performance metrics for the training images, but with validation images it had the lowest performance among all classes. There are two possibilities for this low performance of this class. One is that the classification model was confident in classifying some partially obscured images as non-blurry where the human annotator was not (Figures S12-S13). Second was the low sample size of the blurry image class, which had less than 70 total mid-day images. This likely resulted in the over-fitting of the blurry class on the training images and resulting low performance among validation images. Obtaining more PhenoCam images which are blurry or where the field of view was obscured in some way would be beneficial, and could be obtained from the numerous non-agricultural sites. Regardless, the low accuracy of blurry images had little effect on the final results, since the final classification of any single day is determined by the joint classifications of all surrounding days in the post-processing.

## 5. Conclusion

Monitoring and assessing crop extent and status using a consistent, data-driven approach is essential to meeting the growing demand for food while meeting our sustainability goals in light of climate change. We formulated a workflow using a deep learning model applied to PhenoCam time series to generate a daily crop phenology time series for locations across the continental U.S. The workflow uses a hidden markov model to account for the temporal correlation of daily images. By using images from the full PhenoCam database our model workflow is more resilient since it can accommodate an array of real-world conditions. The resulting outputs offer a ground truth to calibrate and refine existing models for mapping crop status and yield using satellite remote sensing.

## Supplementary Materials

The following are available online at https://www.mdpi.com/2072-4292/1/1/0/s1, Figure S1: Locations of all PhenoCam sites used in the study, Figures S2-S3: Confusion matrices for the classification results, Figures S4-S13: Example classification output from the VGG16 classifier, Figures S14-S15: CNN Model comparison figures, Table S1: Site information for PhenoCams used in this study, Tables S2-S3: Transition matrices for the hidden markove models.

## Author Contributions

Conceptualization, S.T. and D.B.; methodology, S.T.; formal analysis, S.T.; writing—original draft preparation, S.T.; writing—review and editing,S.T. and D.B.; supervision, D.B.; funding acquisition, D.B. All authors have read and agreed to the published version of the manuscript.

## Data Availability Statement

All code and data to reproduce this analysis, as well as the final model predictions, are available in a Zenodo repository (https://doi.org/10.5281/zenodo.5579797).

## Acknowledgments

We thank Andrew Richardson and Koen Hufkens for feedback on our methodology. This research was a contribution from the Long-Term Agroecosystem Research (LTAR) network. LTAR is supported by the United States Department of Agriculture. The authors acknowledge the USDA Agricultural Research Service (ARS) Big Data Initiative and SCINet high performance computing resources (https://scinet.usda.gov) and funding from the Scientific Computing Initiative (SCINet) Postdoctoral Fellow program to support SDT. Any use of trade, firm, or product names is for descriptive purposes only and does not imply endorsement by the U.S. Government. USDA is an equal opportunity provider and employer.

We thank our many collaborators, including site PIs and technicians, for their efforts in support of PhenoCam. The development of PhenoCam Network has been funded by the Northeastern States Research Cooperative, NSF’s Macrosystems Biology program (awards EF-1065029 and EF-1702697), and DOE’s Regional and Global Climate Modeling program (award DE-SC0016011). AmeriFlux is sponsored by the U.S. Department of Energy’s Office of Science.

This research was supported in part by an appointment to the Agricultural Research Service (ARS) Research Participation Program administered by the Oak Ridge Institute for Science and Education (ORISE) through an interagency agreement between the U.S. Department of Energy (DOE) and the U.S. Department of Agriculture (USDA). ORISE is managed by ORAU under DOE contract number DE-SC0014664. All opinions expressed in this paper are the author’s and do not necessarily reflect the policies and views of USDA, DOE, or ORAU/ORISE.

## References

1. Weiss, M.; Jacob, F.; Duveiller, G. Remote sensing for agricultural applications: A meta-review. Remote Sensing of Environment 2020, 236, 111402. doi:10.1016/j.rse.2019.111402.

2. Bégué, A.; Arvor, D.; Bellon, B.; Betbeder, J.; de Abelleyra, D.; Ferraz, R.P.; Lebourgeois, V.; Lelong, C.; Simões, M.; Verón, S.R. Remote sensing and cropping practices: A review. Remote Sensing 2018, 10, 99. doi:10.3390/rs10010099.

3. Gao, F.; Anderson, M.C.; Zhang, X.; Yang, Z.; Alfieri, J.G.; Kustas, W.P.; Mueller, R.; Johnson, D.M.; Prueger, J.H. Toward mapping crop progress at field scales through fusion of Landsat and MODIS imagery. Remote Sensing of Environment 2017, 188, 9–25. doi:10.1016/j.rse.2016.11.004.

4. Gao, F.; Zhang, X. Mapping Crop Phenology in Near Real-Time Using Satellite Remote Sensing: Challenges and Opportunities. Journal of Remote Sensing 2021, 2021, 1–14. doi: 10.34133/2021/8379391.

5. Hufkens, K.; Melaas, E.K.; Mann, M.L.; Foster, T.; Ceballos, F.; Robles, M.; Kramer, B. Monitoring crop phenology using a smartphone based near-surface remote sensing approach. Agricultural and Forest Meteorology 2019, 265, 327–337. doi:10.1016/j.agrformet.2018.11.002.

6. Liu, Y.; Bachofen, C.; Wittwer, R.; Silva Duarte, G.; Sun, Q.; Klaus, V.H.; Buchmann, N. Using PhenoCams to track crop phenology and explain the effects of different cropping systems on yield. Agricultural Systems 2022, 195, 103306. doi:10.1016/j.agsy.2021.103306.

7. Richardson, A.D.; Hufkens, K.; Milliman, T.; Aubrecht, D.M.; Chen, M.; Gray, J.M.; Johnston, M.R.; Keenan, T.F.; Klosterman, S.T.; Kosmala, M.; Melaas, E.K.; Friedl, M.A.; Frolking, S. Tracking vegetation phenology across diverse North American biomes using PhenoCam imagery. Scientific Data 2018, 5, 180028. doi:10.1038/sdata.2018.28.

8. Seyednasrollah, B.; Young, A.M.; Hufkens, K.; Milliman, T.; Friedl, M.A.; Frolking, S.; Richardson, A.D. Tracking vegetation phenology across diverse biomes using Version 2.0 of the PhenoCam Dataset. Scientific data 2019, 6, 222. doi:10.1038/s41597-019-0229-9.

9. Browning, D.M.; Russell, E.S.; Ponce-Campos, G.E.; Kaplan, N.; Richardson, A.D.; Seyednasrollah, B.; Spiegal, S.; Saliendra, N.; Alfieri, J.G.; Baker, J.; Bernacchi, C.; Bestelmeyer, B.T.; Bosch, D.; Boughton, E.H.; Boughton, R.K.; Clark, P.; Flerchinger, G.; Gomez-Casanovas, N.; Goslee, S.; Haddad, N.M.; Hoover, D.; Jaradat, A.; Mauritz, M.; McCarty, G.W.; Miller, G.R.; Sadler, J.; Saha, A.; Scott, R.L.; Suyker, A.; Tweedie, C.; Wood, J.D.; Zhang, X.; Taylor, S.D. Monitoring agroecosystem productivity and phenology at a national scale: A metric assessment framework. Ecological Indicators 2021, 131, 108147. doi:10.1016/j.ecolind.2021.108147.

10. Borowiec, M.L.; Frandsen, P.; Dikow, R.; McKeeken, A.; Valentini, G.; White, A.E. Deep learning as a tool for ecology and evolution. EcoEvoRxiv 2021. doi:10.32942/osf.io/nt3as.

11. Weinstein, B.G. A computer vision for animal ecology. Journal of Animal Ecology 2018, 87, 533–545. doi:10.1111/1365-2656.12780.

12. Norouzzadeh, M.S.; Nguyen, A.; Kosmala, M.; Swanson, A.; Palmer, M.S.; Packer, C.; Clune, J. Automatically identifying, counting, and describing wild animals in camera-trap images with deep learning. Proceedings of the National Academy of Sciences of the United States of America 2018, 115, E5716–E5725. doi:10.1073/pnas.1719367115.

13. Conway, A.M.; Durbach, I.N.; McInnes, A.; Harris, R.N. Frame-by-frame annotation of video recordings using deep neural networks. Ecosphere 2021, 12. doi:10.1002/ecs2.3384.

14. Correia, D.L.; Bouachir, W.; Gervais, D.; Pureswaran, D.; Kneeshaw, D.D.; De Grandpre, L. Leveraging Artificial Intelligence for Large-Scale Plant Phenology Studies from Noisy Time-Lapse Images. IEEE Access 2020, 8, 13151–13160. doi:10.1109/ACCESS.2020.2965462.

15. Kim, T.K.; Kim, S.; Won, M.; Lim, J.H.; Yoon, S.; Jang, K.; Lee, K.H.; Park, Y.D.; Kim, H.S. Utilizing machine learning for detecting flowering in mid-range digital repeat photography. Ecological Modelling 2021, 440, 109419. doi:10.1016/j.ecolmodel.2020.109419.

16. Jones, H.G. What plant is that? Tests of automated image recognition apps for plant identification on plants from the British flora. AoB PLANTS 2020, 12. doi:10.1093/aobpla/plaa052.

17. Ghosal, S.; Blystone, D.; Singh, A.K.; Ganapathysubramanian, B.; Singh, A.; Sarkar, S. An explainable deep machine vision framework for plant stress phenotyping. Proceedings of the National Academy of Sciences 2018, 115, 4613–4618. doi:10.1073/pnas.1716999115.

18. Kosmala, M.; Crall, A.; Cheng, R.; Hufkens, K.; Henderson, S.; Richardson, A. Season Spotter: Using Citizen Science to Validate and Scale Plant Phenology from Near-Surface Remote Sensing. Remote Sensing 2016, 8, 726. doi:10.3390/rs8090726.

19. Song, G.; Wu, S.; Lee, C.K.; Serbin, S.P.; Wolfe, B.T.; Ng, M.K.; Ely, K.S.; Bogonovich, M.; Wang, J.; Lin, Z.; Saleska, S.; Nelson, B.W.; Rogers, A.; Wu, J. Monitoring leaf phenology in moist tropical forests by applying a superpixel-based deep learning method to time-series images of tree canopies. ISPRS Journal of Photogrammetry and Remote Sensing 2022, 183, 19–33. doi: 10.1016/j.isprsjprs.2021.10.023.

20. Cao, M.; Sun, Y.; Jiang, X.; Li, Z.; Xin, Q. Identifying Leaf Phenology of Deciduous Broadleaf Forests from PhenoCam Images Using a Convolutional Neural Network Regression Method. Remote Sensing 2021, 13, 2331. doi:10.3390/rs13122331.

21. Benos, L.; Tagarakis, A.C.; Dolias, G.; Berruto, R.; Kateris, D.; Bochtis, D. Machine Learning in Agriculture: A Comprehensive Updated Review. Sensors 2021, 21, 3758. doi:10.3390/s21113758.

22. Yalcin, H. Plant phenology recognition using deep learning: Deep-Pheno. 2017 6th International Conference on Agro-Geoinformatics, Agro-Geoinformatics 2017 2017. doi:10.1109/Agro-Geoinformatics.2017.8046996.

23. Han, J.; Shi, L.; Yang, Q.; Huang, K.; Zha, Y.; Yu, J. Real-time detection of rice phenology through convolutional neural network using handheld camera images. Precision Agriculture 2021, 22, 154–178. doi:10.1007/s11119-020-09734-2.

24. A.H.M, N.C.; Alkady, K.H.; Jin, H.; Bai, F.; Samal, A.; Ge, Y. A Deep Convolutional Neural Network Based Image Processing Framework for Monitoring the Growth of Soybean Crops. 2021 ASABE Annual International Virtual Meeting, July 12-16, 2021; American Society of Agricultural and Biological Engineers: St. Joseph, MI, 2021; Vol. 2, pp. 754–770. doi:10.13031/aim.202100259.

25. Richardson, A.D. Tracking seasonal rhythms of plants in diverse ecosystems with digital camera imagery. New Phytologist 2019, 222, 1742–1750. doi:10.1111/nph.15591.

26. Aasen, H.; Kirchgessner, N.; Walter, A.; Liebisch, F. PhenoCams for Field Phenotyping: Using Very High Temporal Resolution Digital Repeated Photography to Investigate Interactions of Growth, Phenology, and Harvest Traits. Frontiers in Plant Science 2020, 11. doi: 10.3389/fpls.2020.00593.

27. Barve, V.V.; Brenskelle, L.; Li, D.; Stucky, B.J.; Barve, N.V.; Hantak, M.M.; McLean, B.S.; Paluh, D.J.; Oswald, J.A.; Belitz, M.W.; Folk, R.A.; Guralnick, R.P. Methods for broad-scale plant phenology assessments using citizen scientists’ photographs. Applications in Plant Sciences 2020, 8, 754275. doi:10.1002/aps3.11315.

28. Meier, U. Growth stages of mono-and dicotyledonous plants; Blackwell Wissenschafts-Verlag, 1997.

29. Simonyan, K.; Zisserman, A. Very Deep Convolutional Networks for Large-Scale Image Recognition. arXiv 2014, [arXiv:cs.CV/1409.1556].

30. Chollet, F. Keras: The python deep learning library. Astrophysics Source Code Library 2018, pp. ascl—1806.

31. Milliman, T.; Seyednasrollah, B.; Young, A.M.; Hufkens, K.; Friedl, M.A.; Frolking, S.; Richardson, A.D.; Abraha, M.; Allen, D.W.; Apple, M.; Arain, M.A.; Baker, J.; Baker, J.M.; Bernacchi, C.J.; Bhattacharjee, J.; Blanken, P.; Bosch, D.D.; Boughton, R.; Boughton, E.H.; Brown, R.F.; Browning, D.M.; Brunsell, N.; Burns, S.P.; Cavagna, M.; Chu, H.; Clark, P.E.; Conrad, B.J.; Cre-monese, E.; Debinski, D.; Desai, A.R.; Diaz-Delgado, R.; Duchesne, L.; Dunn, A.L.; Eissenstat, D.M.; El-Madany, T.; Ellum, D.S.S.; Ernest, S.M.; Esposito, A.; Fenstermaker, L.; Flanagan, L.B.; Forsythe, B.; Gallagher, J.; Gianelle, D.; Griffis, T.; Groffman, P.; Gu, L.; Guillemot, J.; Halpin, M.; Hanson, P.J.; Hemming, D.; Hove, A.A.; Humphreys, E.R.; Jaimes-Hernandez, A.; Jaradat, A.A.; Johnson, J.; Keel, E.; Kelly, V.R.; Kirchner, J.W.; Kirchner, P.B.; Knapp, M.; Krassovski, M.; Langvall, O.; Lanthier, G.; Maire, G.; Magliulo, E.; Martin, T.A.; McNeil, B.; Meyer, G.A.; Migliavacca, M.; Mohanty, B.P.; Moore, C.E.; Mudd, R.; Munger, J.W.; Murrell, Z.E.; Nesic, Z.; Neufeld, H.S.; Oechel, W.; Oishi, A.C.; Oswald, W.W.; Perkins, T.D.; Reba, M.L.; Rundquist, B.; Runkle, B.R.; Russell, E.S.; Sadler, E.J.; Saha, A.; Saliendra, N.Z.; Schmalbeck, L.; Schwartz, M.D.; Scott, R.L.; Smith, E.M.; Sonnentag, O.; Stoy, P.; Strachan, S.; Suvocarev, K.; Thom, J.E.; Thomas, R.Q.; Den berg, A.K.; Vargas, R.; Vogel, C.S.; Walker, J.J.; Webb, N.; Wetzel, P.; Weyers, S.; Whipple, A.V.; Whitham, T.G.; Wohlfahrt, G.; Wood, J.D.; Yang, J.; Yang, X.; Yenni, G.; Zhang, Y.; Zhang, Q.; Zona, D.; Baldocchi, D.; Verfaillie, J. PhenoCam Dataset v2.0: Digital Camera Imagery from the PhenoCam Network, 2000-2018, 2019. doi:10.3334/ORNLDAAC/1689.

32. Esmael, B.; Arnaout, A.; Fruhwirth, R.K.; Thonhauser, G. Improving time series classification using Hidden Markov Models. 2012 12th International Conference on Hybrid Intelligent Systems (HIS). IEEE, 2012, pp. 502–507. doi:10.1109/HIS.2012.6421385.

33. Wehmann, A.; Liu, D. A spatial–temporal contextual Markovian kernel method for multitemporal land cover mapping. ISPRS Journal of Photogrammetry and Remote Sensing 2015, 107, 77–89. doi:10.1016/j.isprsjprs.2015.04.009.

34. Abercrombie, S.P.; Friedl, M.A. Improving the Consistency of Multitemporal Land Cover Maps Using a Hidden Markov Model. IEEE Transactions on Geoscience and Remote Sensing 2016, 54, 703–713. doi:10.1109/TGRS.2015.2463689.

35. Abadi, M.; Agarwal, A.; Barham, P.; Brevdo, E.; Chen, Z.; Citro, C.; Corrado, G.S.; Davis, A.; Dean, J.; Devin, M.; Ghemawat, S.; Goodfellow, I.; Harp, A.; Irving, G.; Isard, M.; Jia, Y.; Jozefowicz, R.; Kaiser, L.; Kudlur, M.; Levenberg, J.; Mané, D.; Monga, R.; Moore, S.; Murray, D.; Olah, C.; Schuster, M.; Shlens, J.; Steiner, B.; Sutskever, I.; Talwar, K.; Tucker, P.; Vanhoucke, V.; Vasudevan, V.; Viégas, F.; Vinyals, O.; Warden, P.; Wattenberg, M.; Wicke, M.; Yu, Y.; Zheng, X. TensorFlow: Large-Scale Machine Learning on Heterogeneous Systems. arXiv 2016, [arXiv:cs.DC/1603.04467].

36. McKinney, W. Data Structures for Statistical Computing in Python. Proceedings of the 9th Python in Science Conference; SciPy: Austin, Texas, USA, 2010; pp. 51–56.

37. Harris, C.R.; Millman, K.J.; van der Walt, S.J.; Gommers, R.; Virtanen, P.; Cournapeau, D.; Wieser, E.; Taylor, J.; Berg, S.; Smith, N.J.; Kern, R.; Picus, M.; Hoyer, S.; van Kerkwijk, M.H.; Brett, M.; Haldane, A.; del Río, J.F.; Wiebe, M.; Peterson, P.; Gérard-Marchant, P.; Sheppard, K.; Reddy, T.; Weckesser, W.; Abbasi, H.; Gohlke, C.; Oliphant, T.E. Array programming with NumPy. Nature 2020, 585, 357–362. doi:10.1038/s41586-020-2649-2.

38. Schreiber, J. Pomegranate: fast and flexible probabilistic modeling in python. Journal of Machine Learning Research 2018, 18, 1–6.

39. Python Software Foundation. Python Language Reference Manual, version 3.6. url- http://www.python.org, 2003.

40. R Core Team. R: a language and environment for statistical computing, 2017.

41. Zeileis, A.; Grothendieck, G. zoo: S3 Infrastructure for Regular and Irregular Time Series. Journal of Statistical Software 2005, 14. doi:10.18637/jss.v014.i06.

42. Wickham, H.; Averick, M.; Bryan, J.; Chang, W.; McGowan, L.; François, R.; Grolemund, G.; Hayes, A.; Henry, L.; Hester, J.; Kuhn, M.; Pedersen, T.; Miller, E.; Bache, S.; Müller, K.; Ooms, J.; Robinson, D.; Seidel, D.; Spinu, V.; Takahashi, K.; Vaughan, D.; Wilke, C.; Woo, K.; Yutani, H. Welcome to the Tidyverse. Journal of Open Source Software 2019, 4, 1686. doi:10.21105/joss.01686.

43. Wickham, H. ggplot2: Elegant Graphics for Data Analysis; Springer-Verlag New York, 2016.

44. Taylor, S.D. Analysis code for: Deep learning models for identifying crop and field attributes from near surface cameras., 2021. doi:10.5281/zenodo.5579797.

45. USDA-NASS. The Yield Forecasting Program of NASS. Report SMB 12-01. Technical report, USDA National Agricultural Statistics Service, 2012.

46. Lombardozzi, D.L.; Lu, Y.; Lawrence, P.J.; Lawrence, D.M.; Swenson, S.; Oleson, K.W.; Wieder, W. R.; Ainsworth, E.A. Simulating agriculture in the Community Land Model version 5. Journal of Geophysical Research: Biogeosciences 2020, pp. 0–3. doi:10.1029/2019JG005529.

47. Richardson, A.D.; Hufkens, K.; Milliman, T.; Frolking, S. Intercomparison of phenological transition dates derived from the PhenoCam Dataset V1.0 and MODIS satellite remote sensing. Scientific Reports 2018, 8, 5679. doi:10.1038/s41598-018-23804-6.

48. Sakamoto, T.; Wardlow, B.D.; Gitelson, A.A.; Verma, S.B.; Suyker, A.E.; Arkebauer, T.J. A Two-Step Filtering approach for detecting maize and soybean phenology with time-series MODIS data. Remote Sensing of Environment 2010, 114, 2146–2159. doi:10.1016/j.rse.2010.04.019.

49. Fine, S.; Singer, Y.; Tishby, N. The hierarchical hidden Markov model: Analysis and applications. Machine learning 1998, 32, 41–62. doi:10.1023/A:1007469218079.

50. Lea, C.; Reiter, A.; Vidal, R.; Hager, G.D. Segmental Spatiotemporal CNNs for Fine-Grained Action Segmentation. In Computer Vision – ECCV 2016; B., L.; J., M.; N., S.; M., W., Eds.; Springer, Cham, 2016; Vol. 9907, pp. 36–52, [1602.02995]. doi:10.1007/978-3-319-46487-9_3.

