## Supplemental Info for "Classification of daily crop phenology in PhenoCams using deep learning and hidden markov models"

Supplemental Images S1-S15
Supplemental Table S1-S3

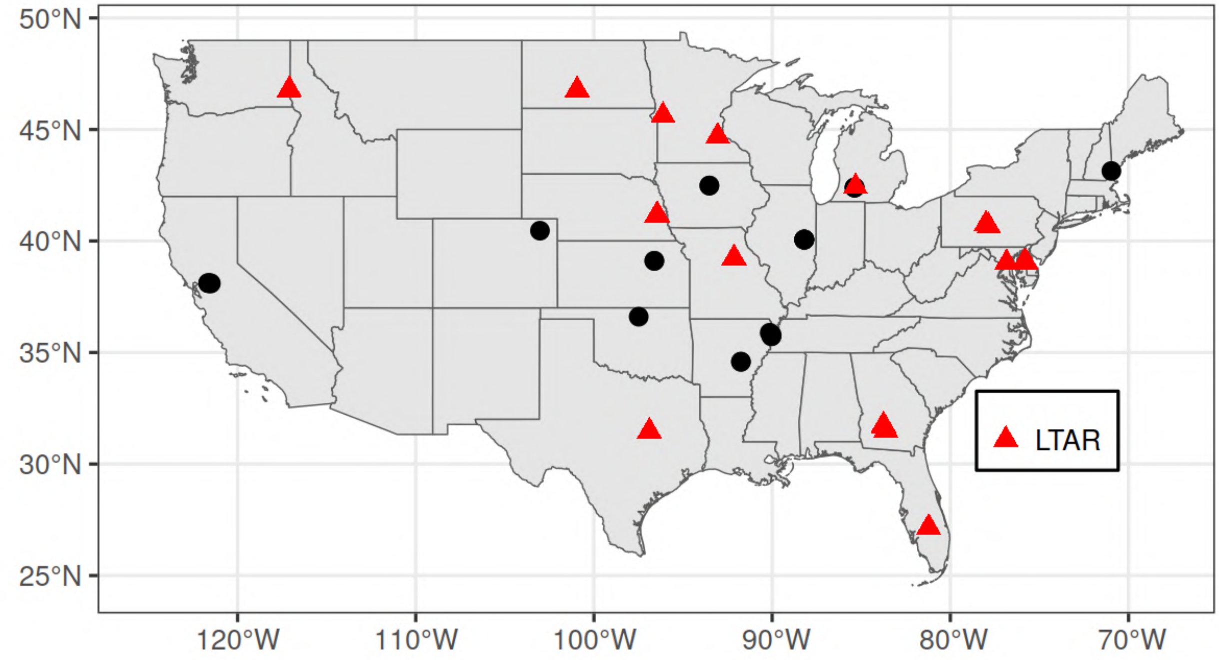


Figure S1: Location of all PhenoCam sites used in the study. LTAR sites (red) are part of the Long-Term Agroecosystem Research Network in addition to the PhenoCam network. Three additional cameras are located in Italy (borgocioffinorth, borgocioffisourth) and China (jurong).

| | phenocam_name | ltar_site | lat | lon | roi_type | roi_id | first_date | last_date | site_years |
|---|---|---|---|---|---|---|---|---|---|
| 1 | archboldwet | yes | 27.16557 | -81.21613 | AG | 1000 | 2016-10-31 | 2021-07-07 | 2.5 |
| 2 | arscolesnorth | no | 42.48840 | -93.52250 | AG | 1000 | 2018-08-21 | 2021-07-07 | 2.1 |
| 3 | arscolessouth | no | 42.48160 | -93.52350 | AG | 1000 | 2018-08-18 | 2021-07-07 | 2.3 |
| 4 | arsgacp1 | yes | 31.51090 | -83.61790 | AG | 1000 | 2016-05-10 | 2021-07-07 | 5.2 |
| 5 | arsgacp3 | yes | 31.75630 | -83.75160 | AG | 1000 | 2018-04-19 | 2021-07-04 | 3.1 |
| 6 | arsgacp4 | yes | 31.70930 | -83.72870 | AG | 1000 | 2018-06-22 | 2021-07-07 | 2.1 |
| 7 | arsltarmdcr | yes | 39.05870 | -75.85130 | AG | 1000 | 2017-05-11 | 2021-07-07 | 4.2 |
| 8 | arsltarmdcrprcw | yes | 39.06670 | -75.76090 | AG | 1000 | 2019-02-16 | 2021-07-07 | 2.4 |
| 9 | arsltarmdcrresw | yes | 39.05490 | -75.75320 | AG | 1000 | 2019-03-18 | 2021-07-07 | 2.3 |
| 10 | arsltarucbec1 | yes | 40.75370 | -78.00570 | AG | 1000 | 2019-05-17 | 2021-07-07 | 2.1 |
| 11 | arsmorris1 | yes | 45.61670 | -96.12690 | AG | 1000 | 2017-07-25 | 2021-07-04 | 4.0 |
| 12 | arsmorris2 | yes | 45.62700 | -96.12700 | AG | 1000 | 2017-07-25 | 2021-07-04 | 3.9 |
| 13 | arsope3ltar | yes | 39.03090 | -76.84420 | AG | 1000 | 2017-04-11 | 2021-07-04 | 4.2 |
| 14 | borgocioffinorth | no | 40.52370 | 14.95740 | AG | 1000 | 2017-02-16 | 2021-07-01 | 4.0 |
| 15 | borgocioffisouth | no | 40.52370 | 14.95740 | AG | 1000 | 2017-03-15 | 2021-07-01 | 4.0 |
| 16 | bouldinalfalfa | no | 38.09850 | -121.49930 | AG | 1000 | 2016-11-03 | 2021-06-22 | 4.3 |
| 17 | bouldincorn | no | 38.10900 | -121.53500 | AG | 1000 | 2017-07-13 | 2021-06-22 | 3.7 |
| 18 | burdettericea | no | 35.80890 | -90.03271 | AG | 1000 | 2015-06-19 | 2018-05-17 | 2.1 |
| 19 | cafboydnorthltar01 | yes | 46.75510 | -117.12605 | AG | 1000 | 2017-09-14 | 2021-07-07 | 3.4 |
| 20 | cafcookeastltar01 | yes | 46.78152 | -117.08205 | AG | 1000 | 2017-05-08 | 2021-07-07 | 4.1 |
| 21 | cafcookwestltar01 | yes | 46.78404 | -117.09083 | AG | 1000 | 2017-06-25 | 2021-07-07 | 3.7 |
| 22 | goodwater | yes | 39.22848 | -92.11936 | AG | 1000 | 2015-09-26 | 2021-06-19 | 5.7 |
| 23 | goodwaterbau | yes | 39.23115 | -92.15216 | AG | 1000 | 2018-10-02 | 2021-07-07 | 2.8 |
| 24 | hawbeckereddy | yes | 40.66080 | -77.84885 | AG | 1000 | 2015-09-23 | 2019-05-14 | 3.4 |
| 25 | humnokericea | no | 34.58519 | -91.75168 | AG | 1000 | 2015-06-25 | 2021-07-07 | 5.2 |
| 26 | humnokericec | no | 34.58885 | -91.75167 | AG | 1000 | 2015-06-25 | 2021-07-07 | 5.5 |
| 27 | jurong | no | 31.80680 | 119.21730 | AG | 1000 | 2017-10-23 | 2021-07-07 | 3.6 |
| 28 | kelloggcorn | yes | 42.43754 | -85.32255 | AG | 1000 | 2014-05-23 | 2019-10-05 | 4.9 |
| 29 | kelloggcornsoy | no | 42.39601 | -85.37529 | AG | 1000 | 2015-07-16 | 2020-06-15 | 3.4 |
| 30 | kelloggcornsoy2 | no | 42.39576 | -85.37425 | AG | 1000 | 2015-07-16 | 2021-07-07 | 5.1 |
| 31 | kelloggnativegrass | no | 42.39576 | -85.37456 | AG | 1000 | 2015-08-18 | 2018-03-18 | 2.5 |
| 32 | mandanh5 | yes | 46.77542 | -100.95109 | AG | 1000 | 2015-09-17 | 2021-07-07 | 5.8 |
| 33 | mandani2 | yes | 46.76140 | -100.92570 | AG | 1000 | 2016-04-22 | 2021-07-07 | 5.1 |
| 34 | manilacotton | no | 35.88720 | -90.13710 | AG | 1000 | 2016-06-21 | 2021-01-14 | 4.5 |
| 35 | mead1 | yes | 41.16510 | -96.47660 | AG | 1000 | 2016-07-12 | 2021-07-07 | 5.0 |
| 36 | mead2 | yes | 41.16490 | -96.47010 | AG | 1000 | 2016-07-12 | 2021-07-07 | 5.0 |
| 37 | mead3 | yes | 41.17970 | -96.43970 | AG | 1000 | 2016-07-12 | 2021-07-07 | 5.0 |
| 38 | moorefields | no | 43.13830 | -70.96090 | AG | 1000 | 2016-07-06 | 2018-11-13 | 2.3 |
| 39 | NEON.D06.KONA.DP1.00033 | no | 39.11045 | -96.61293 | AG | 1000 | 2016-05-07 | 2021-07-07 | 5.0 |
| 40 | NEON.D06.KONA.DP1.00042 | no | 39.11045 | -96.61293 | AG | 1000 | 2016-05-07 | 2021-07-07 | 5.0 |
| 41 | NEON.D10.STER.DP1.00033 | no | 40.46189 | -103.02929 | AG | 1000 | 2016-12-18 | 2021-07-07 | 4.5 |
| 42 | rosemountcons | yes | 44.69460 | -93.05780 | AG | 1000 | 2017-05-04 | 2020-12-15 | 3.6 |
| 43 | rosemountconv | yes | 44.69100 | -93.05760 | AG | 1000 | 2017-05-19 | 2020-12-21 | 3.6 |
| 44 | southerngreatplains | no | 36.60580 | -97.48880 | AG | 1000 | 2012-05-16 | 2021-07-04 | 8.8 |
| 45 | twitchell | no | 38.10873 | -121.65302 | AG | 1000 | 2011-11-16 | 2017-04-05 | 4.0 |
| 46 | twitchellalfalfa | no | 38.11542 | -121.64667 | AG | 1000 | 2013-05-23 | 2016-09-22 | 2.7 |
| 47 | tworfaa | yes | 31.47770 | -96.88820 | AG | 1000 | 2018-04-20 | 2021-07-07 | 3.2 |
| 48 | uiefmaize | no | 40.06282 | -88.19613 | AG | 1000 | 2008-11-06 | 2017-12-31 | 9.1 |
| 49 | uiefmaize2 | no | 40.06280 | -88.19610 | AG | 1000 | 2018-08-30 | 2021-07-04 | 2.9 |
| 50 | uiefmiscanthus | no | 40.06281 | -88.19843 | AG | 1000 | 2008-11-12 | 2018-04-29 | 9.2 |
| 51 | uiefmiscanthus | no | 40.06281 | -88.19843 | AG | 2000 | 2018-05-02 | 2021-07-04 | 2.9 |
| 52 | uiefswitchgrass | no | 40.06465 | -88.19606 | AG | 1000 | 2008-10-22 | 2018-12-31 | 9.4 |
| 53 | uiefswitchgrass2 | no | 40.06370 | -88.19730 | AG | 1000 | 2018-08-30 | 2021-07-04 | 2.9 |
| 54 | usof5 | no | 35.72970 | -90.04060 | AG | 1000 | 2018-05-20 | 2021-01-14 | 2.3 |
| 55 | usof6 | no | 35.73330 | -90.04030 | AG | 1000 | 2018-05-20 | 2021-01-14 | 2.5 |

Table S1: Site information for PhenoCams used in this study.

| | vegetation | residue | soil | snow | water |
|---|---|---|---|---|---|
| vegetation | 0.95 | 0.0125 | 0.0125 | 0.0125 | 0.0125 |
| residue | 0.0125 | 0.95 | 0.0125 | 0.0125 | 0.0125 |
| soil | 0.0167 | 0 | 0.95 | 0.0167 | 0.0167 |
| snow | 0.0125 | 0.0125 | 0.0125 | 0.95 | 0.0125 |
| water | 0.0125 | 0.0125 | 0.0125 | 0.0125 | 0.95 |

Table S2: Transition probabilities for the Dominant Cover hidden markov model. Rows indicate the state of the current timestep in the latent time series. Columns indicate the potential state of the next timestep (i.e. the next day) and the associated probability of that occurring. For example, transitioning from soil to residue has a probability of 0, while moving from residue to soil has a probability of 0.0125.

| | emergence | growth | flowers | senescing | senesced | no_crop |
|---|---|---|---|---|---|---|
| emergence | 0.9 | 0.05 | 0 | 0 | 0 | 0.05 |
| growth | 0 | 0.9 | 0.025 | 0.025 | 0 | 0.05 |
| flowers | 0 | 0 | 0.9 | 0.05 | 0 | 0.05 |
| senescing | 0 | 0.01 | 0 | 0.9 | 0.045 | 0.045 |
| senesced | 0 | 0.01 | 0 | 0 | 0.9 | 0.09 |
| no_crop | 0.1 | 0 | 0 | 0 | 0 | 0.9 |

Table S3: Transition probabilities for the Crop Status hidden markov model. The structure is the same as Table S2. For example, transitioning from senesced to growth has a probability of 0.01, and transiting from growth to senesced has a probability of 0.

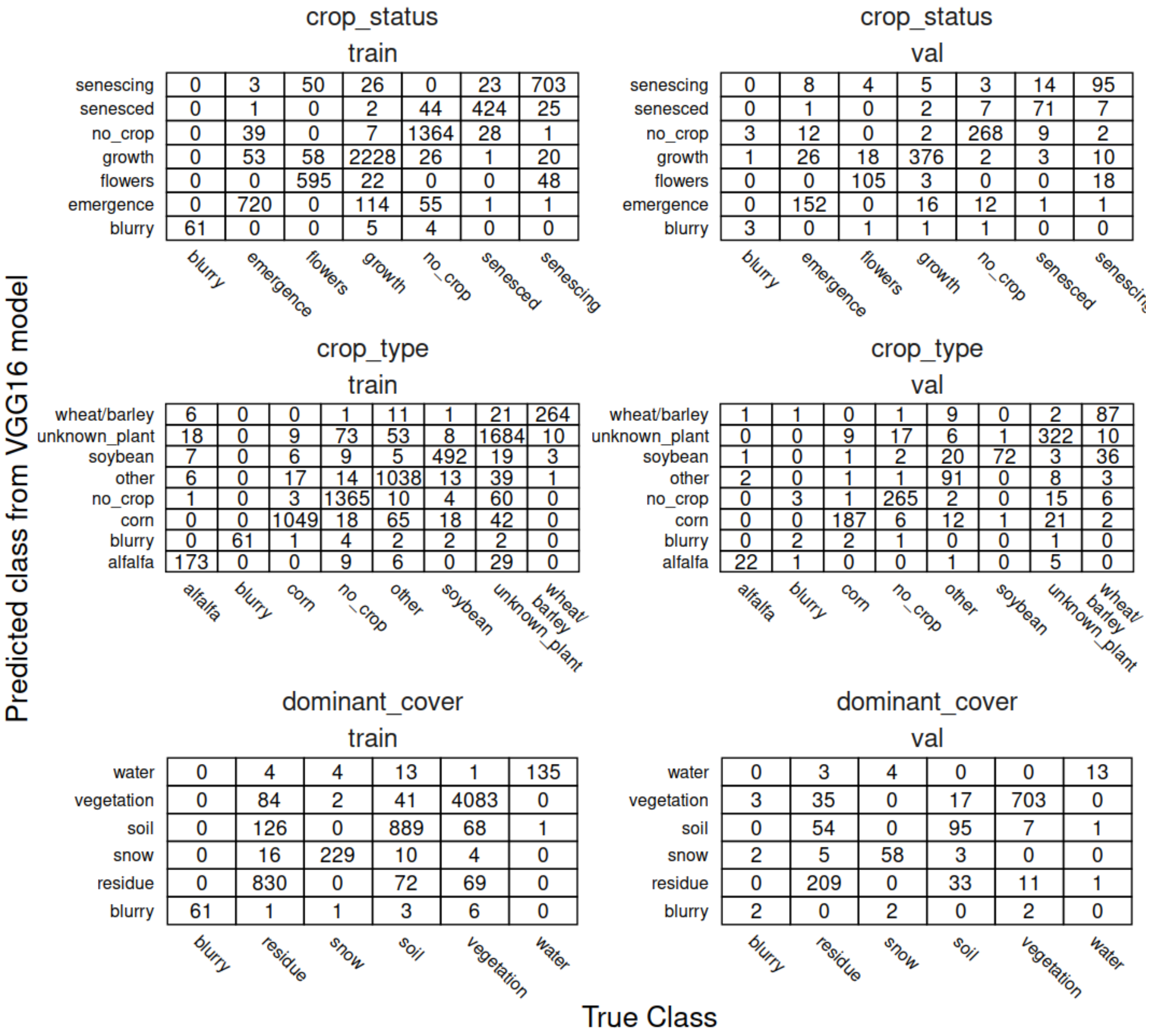



**crop_status — train** (rows: predicted class; columns: true class)

| Predicted \ True | blurry | emergence | flowers | growth | no_crop | senesced | senescing |
|---|---|---|---|---|---|---|---|
| senescing | 0 | 3 | 50 | 26 | 0 | 23 | 703 |
| senesced | 0 | 1 | 0 | 2 | 44 | 424 | 25 |
| no_crop | 0 | 39 | 0 | 7 | 1364 | 28 | 1 |
| growth | 0 | 53 | 58 | 2228 | 26 | 1 | 20 |
| flowers | 0 | 0 | 595 | 22 | 0 | 0 | 48 |
| emergence | 0 | 720 | 0 | 114 | 55 | 1 | 1 |
| blurry | 61 | 0 | 0 | 5 | 4 | 0 | 0 |

**crop_status — val**

| Predicted \ True | blurry | emergence | flowers | growth | no_crop | senesced | senescing |
|---|---|---|---|---|---|---|---|
| senescing | 0 | 8 | 4 | 5 | 3 | 14 | 95 |
| senesced | 0 | 1 | 0 | 2 | 7 | 71 | 7 |
| no_crop | 3 | 12 | 0 | 2 | 268 | 9 | 2 |
| growth | 1 | 26 | 18 | 376 | 2 | 3 | 10 |
| flowers | 0 | 0 | 105 | 3 | 0 | 0 | 18 |
| emergence | 0 | 152 | 0 | 16 | 12 | 1 | 1 |
| blurry | 3 | 0 | 1 | 1 | 1 | 0 | 0 |

**crop_type — train**

| Predicted \ True | alfalfa | blurry | corn | no_crop | other | soybean | unknown_plant | wheat/barley |
|---|---|---|---|---|---|---|---|---|
| wheat/barley | 6 | 0 | 0 | 1 | 11 | 1 | 21 | 264 |
| unknown_plant | 18 | 0 | 9 | 73 | 53 | 8 | 1684 | 10 |
| soybean | 7 | 0 | 6 | 9 | 5 | 492 | 19 | 3 |
| other | 6 | 0 | 17 | 14 | 1038 | 13 | 39 | 1 |
| no_crop | 1 | 0 | 3 | 1365 | 10 | 4 | 60 | 0 |
| corn | 0 | 0 | 1049 | 18 | 65 | 18 | 42 | 0 |
| blurry | 0 | 61 | 1 | 4 | 2 | 2 | 2 | 0 |
| alfalfa | 173 | 0 | 0 | 9 | 6 | 0 | 29 | 0 |

**crop_type — val**

| Predicted \ True | alfalfa | blurry | corn | no_crop | other | soybean | unknown_plant | wheat/barley |
|---|---|---|---|---|---|---|---|---|
| wheat/barley | 1 | 1 | 0 | 1 | 9 | 0 | 2 | 87 |
| unknown_plant | 0 | 0 | 9 | 17 | 6 | 1 | 322 | 10 |
| soybean | 1 | 0 | 1 | 2 | 20 | 72 | 3 | 36 |
| other | 2 | 0 | 1 | 1 | 91 | 0 | 8 | 3 |
| no_crop | 0 | 3 | 1 | 265 | 2 | 0 | 15 | 6 |
| corn | 0 | 0 | 187 | 6 | 12 | 1 | 21 | 2 |
| blurry | 0 | 2 | 2 | 1 | 0 | 0 | 1 | 0 |
| alfalfa | 22 | 1 | 0 | 0 | 1 | 0 | 5 | 0 |

**dominant_cover — train**

| Predicted \ True | blurry | residue | snow | soil | vegetation | water |
|---|---|---|---|---|---|---|
| water | 0 | 4 | 4 | 13 | 1 | 135 |
| vegetation | 0 | 84 | 2 | 41 | 4083 | 0 |
| soil | 0 | 126 | 0 | 889 | 68 | 1 |
| snow | 0 | 16 | 229 | 10 | 4 | 0 |
| residue | 0 | 830 | 0 | 72 | 69 | 0 |
| blurry | 61 | 1 | 1 | 3 | 6 | 0 |

**dominant_cover — val**

| Predicted \ True | blurry | residue | snow | soil | vegetation | water |
|---|---|---|---|---|---|---|
| water | 0 | 3 | 4 | 0 | 0 | 13 |
| vegetation | 3 | 35 | 0 | 17 | 703 | 0 |
| soil | 0 | 54 | 0 | 95 | 7 | 1 |
| snow | 2 | 5 | 58 | 3 | 0 | 0 |
| residue | 0 | 209 | 0 | 33 | 11 | 1 |
| blurry | 2 | 0 | 2 | 0 | 2 | 0 |

Figure S2: Confusion matrix for the initial model classification. Train represents the model training data while val represents the held-out validation data. Total counts represent the original 8,015 annotated images.

crop_status
train

| | emergence | flowers | growth | no_crop | senesced | senescing |
|---|---|---|---|---|---|---|
| senescing | 0 | 44 | 24 | 14 | 23 | 650 |
| senesced | 0 | 0 | 2 | 38 | 405 | 26 |
| no_crop | 40 | 0 | 6 | 1217 | 32 | 1 |
| growth | 49 | 49 | 2087 | 40 | 0 | 21 |
| flowers | 0 | 548 | 20 | 0 | 0 | 40 |
| emergence | 710 | 0 | 93 | 47 | 3 | 0 |

crop_status
val

| | emergence | flowers | growth | no_crop | senesced | senescin |
|---|---|---|---|---|---|---|
| senescing | 6 | 7 | 3 | 4 | 15 | 92 |
| senesced | 0 | 0 | 1 | 8 | 70 | 2 |
| no_crop | 12 | 0 | 0 | 243 | 8 | 1 |
| growth | 27 | 15 | 360 | 3 | 3 | 8 |
| flowers | 0 | 101 | 5 | 0 | 0 | 20 |
| emergence | 151 | 0 | 14 | 7 | 0 | 0 |

crop_type
train

| | alfalfa | corn | no_crop | other | soybean | wheat/barley |
|---|---|---|---|---|---|---|
| wheat/barley | 5 | 0 | 25 | 0 | 0 | 265 |
| soybean | 18 | 0 | 20 | 9 | 502 | 2 |
| other | 19 | 0 | 37 | 1035 | 8 | 0 |
| no_crop | 1 | 4 | 1217 | 9 | 3 | 1 |
| corn | 0 | 1014 | 12 | 34 | 0 | 2 |
| alfalfa | 139 | 0 | 10 | 0 | 0 | 0 |

crop_type
val

| | alfalfa | corn | no_crop | other | soybean | wheat/barley |
|---|---|---|---|---|---|---|
| wheat/barley | 1 | 0 | 2 | 0 | 0 | 130 |
| soybean | 1 | 0 | 3 | 24 | 66 | 0 |
| other | 3 | 0 | 2 | 93 | 0 | 0 |
| no_crop | 0 | 1 | 243 | 1 | 0 | 5 |
| corn | 0 | 195 | 10 | 13 | 0 | 7 |
| alfalfa | 19 | 0 | 1 | 0 | 0 | 0 |

dominant_cover
train

| | residue | snow | soil | vegetation | water |
|---|---|---|---|---|---|
| water | 2 | 1 | 6 | 1 | 118 |
| vegetation | 71 | 4 | 26 | 3822 | 0 |
| soil | 100 | 2 | 871 | 42 | 8 |
| snow | 14 | 205 | 6 | 8 | 0 |
| residue | 833 | 5 | 51 | 72 | 0 |

dominant_cover
val

| | residue | snow | soil | vegetation | water |
|---|---|---|---|---|---|
| water | 1 | 2 | 1 | 0 | 13 |
| vegetation | 36 | 0 | 12 | 669 | 0 |
| soil | 33 | 0 | 92 | 5 | 0 |
| snow | 7 | 53 | 3 | 0 | 0 |
| residue | 222 | 2 | 30 | 13 | 1 |

Predicted class after post-processing

True Class

Figure S3: Confusion matrix for the model classifications after post-processing. Train represents the model training data while val represents the held-out validation data. Total counts represent the original 8,015 annotated images minus the blurry and unknown plant classes. See the main text for details.

Figures S4-S13: These figures represent single images classified by the VGG16 classifier across three categories. The heading indicates the PhenoCam site name, the image acquisition date, and whether it was part of the training dataset (train) or validation dataset (val). The bar graphs indicate the probabilities, as assigned by the classifier, of the image belonging to the respective class. The final classification in each category is shaded in green. The “Human Annotation” text indicates which class the image was originally annotated as. Note results in these images are from the initial VGG16 classification model and have *not* been transformed via any of the post-processing steps described in the main text.

The images in Figures S4-S13 are licensed under a CC-BY license with the following attribution:

Milliman, T., B. Seyednasrollah, A.M. Young, K. Hufkens, M.A. Friedl, S. Frolking, A.D. Richardson, M. Abraha, D.W. Allen, M. Apple, M.A. Arain, J.M. Baker, D. Baldocchi, C.J. Bernacchi, J. Bhattacharjee, P. Blanken, D.D. Bosch, R. Boughton, E.H. Boughton, R.F. Brown, D.M. Browning, N. Brunsell, S.P. Burns, M. Cavagna, H. Chu, P.E. Clark, B.J. Conrad, E. Cremonese, D. Debinski, A.R. Desai, R. Diaz-Delgado, L. Duchesne, A.L. Dunn, D.M. Eissenstat, T. El-Madany, D.S.S. Ellum, S.M. Ernest, A. Esposito, L. Fenstermaker, L.B. Flanagan, B. Forsythe, J. Gallagher, D. Gianelle, T. Griffis, P. Groffman, L. Gu, J. Guillemot, M. Halpin, P.J. Hanson, D. Hemming, A.A. Hove, E.R. Humphreys, A. Jaimes-Hernandez, A.A. Jaradat, J. Johnson, E. Keel, V.R. Kelly, J.W. Kirchner, P.B. Kirchner, M. Knapp, M. Krassovski, O. Langvall, G. Lanthier, G.l. Maire, E. Magliulo, T.A. Martin, B. McNeil, G.A. Meyer, M. Migliavacca, B.P. Mohanty, C.E. Moore, R. Mudd, J.W. Munger, Z.E. Murrell, Z. Nesic, H.S. Neufeld, W. Oechel, A.C. Oishi, W.W. Oswald, T.D. Perkins, M.L. Reba, B. Rundquist, B.R. Runkle, E.S. Russell, E.J. Sadler, A. Saha, N.Z. Saliendra, L. Schmalbeck, M.D. Schwartz, R.L. Scott, E.M. Smith, O. Sonnentag, P. Stoy, S. Strachan, K. Suvocarev, J.E. Thom, R.Q. Thomas, A.K. Van den berg, R. Vargas, J. Verfaillie, C.S. Vogel, J.J. Walker, N. Webb, P. Wetzel, S. Weyers, A.V. Whipple, T.G. Whitham, G. Wohlfahrt, J.D. Wood, J. Yang, X. Yang, G. Yenni, Y. Zhang, Q. Zhang, and D. Zona. 2019. PhenoCam Dataset v2.0: Digital Camera Imagery from the PhenoCam Network, 2000-2018. ORNL DAAC, Oak Ridge, Tennessee, USA. https://doi.org/10.3334/ORNLDAAC/1689

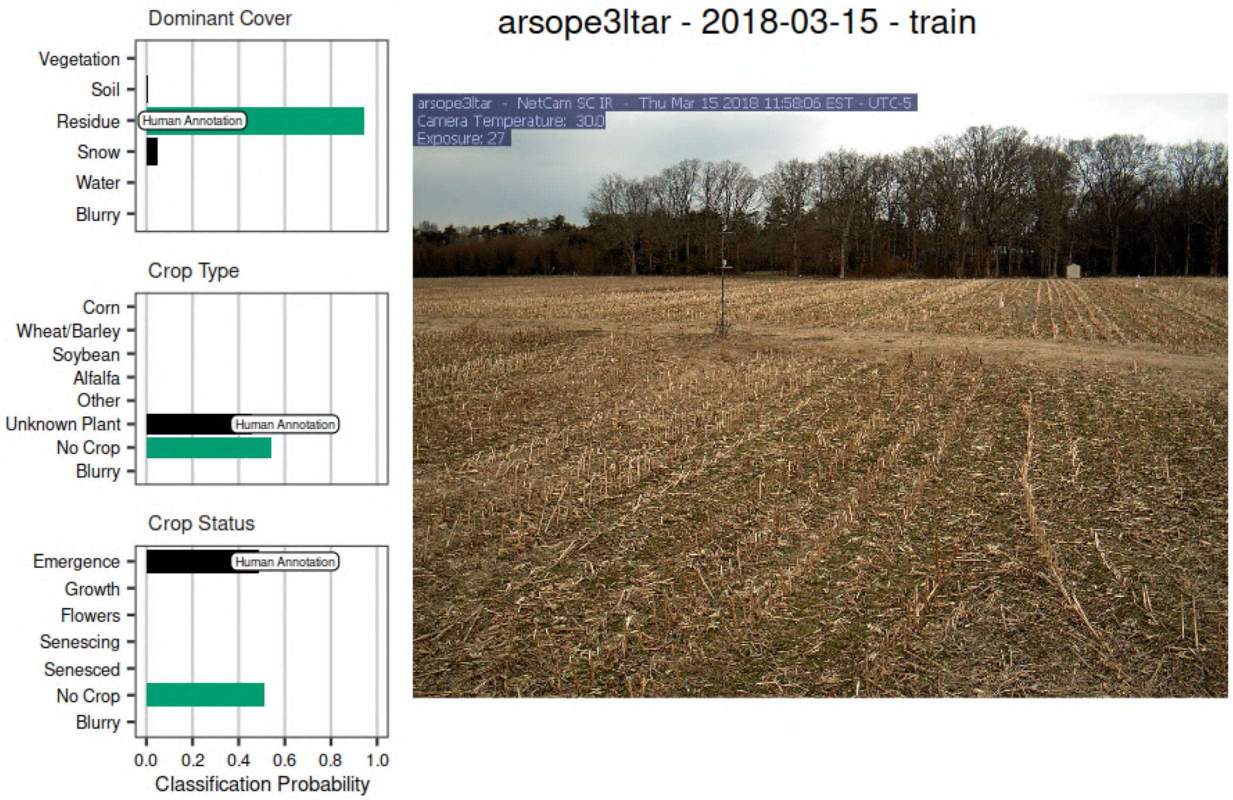


Figure S4

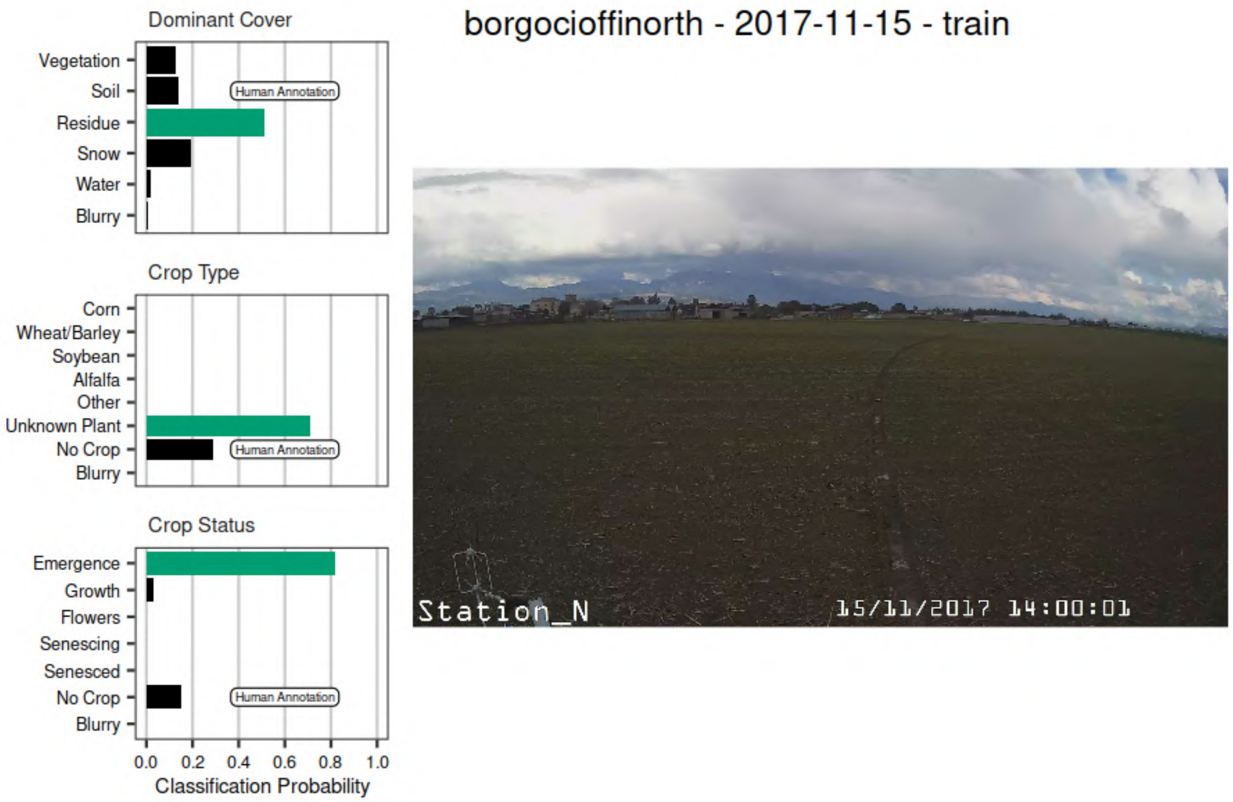


Figure S5

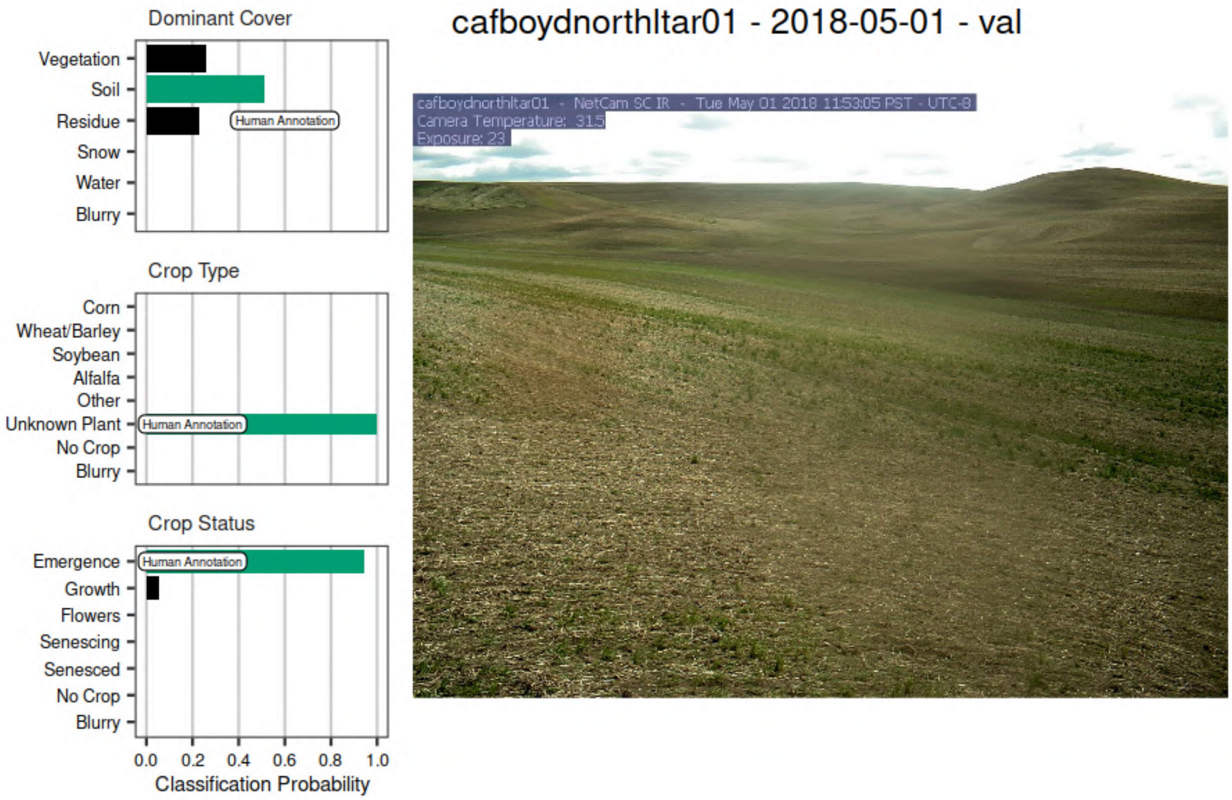


Figure S6

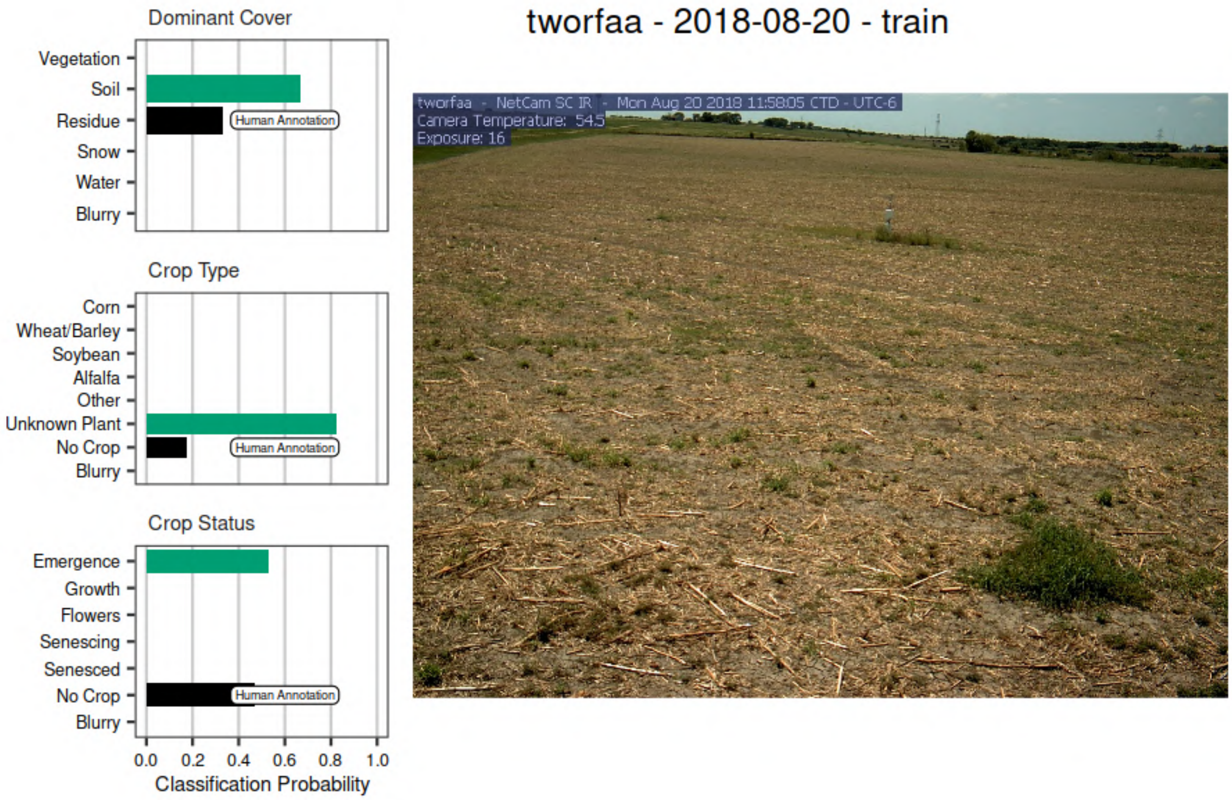


Figure S7

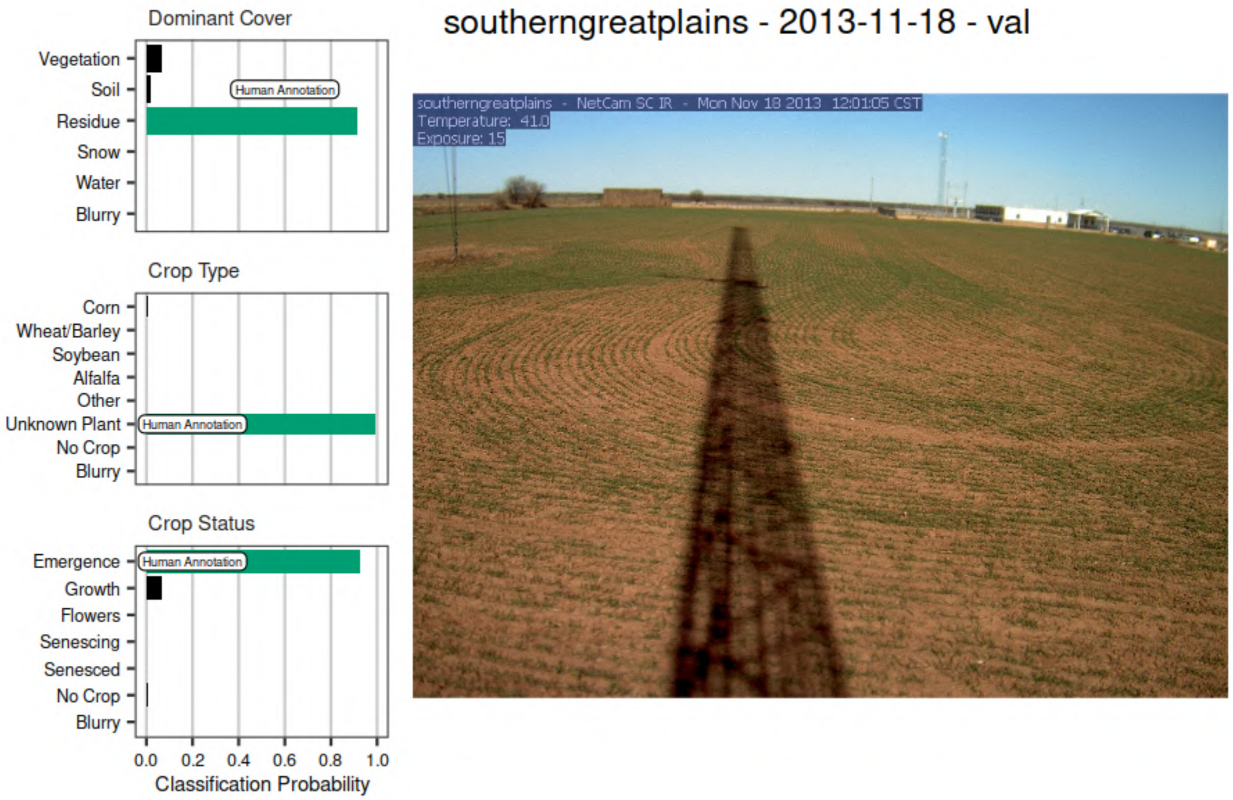


Figure S8

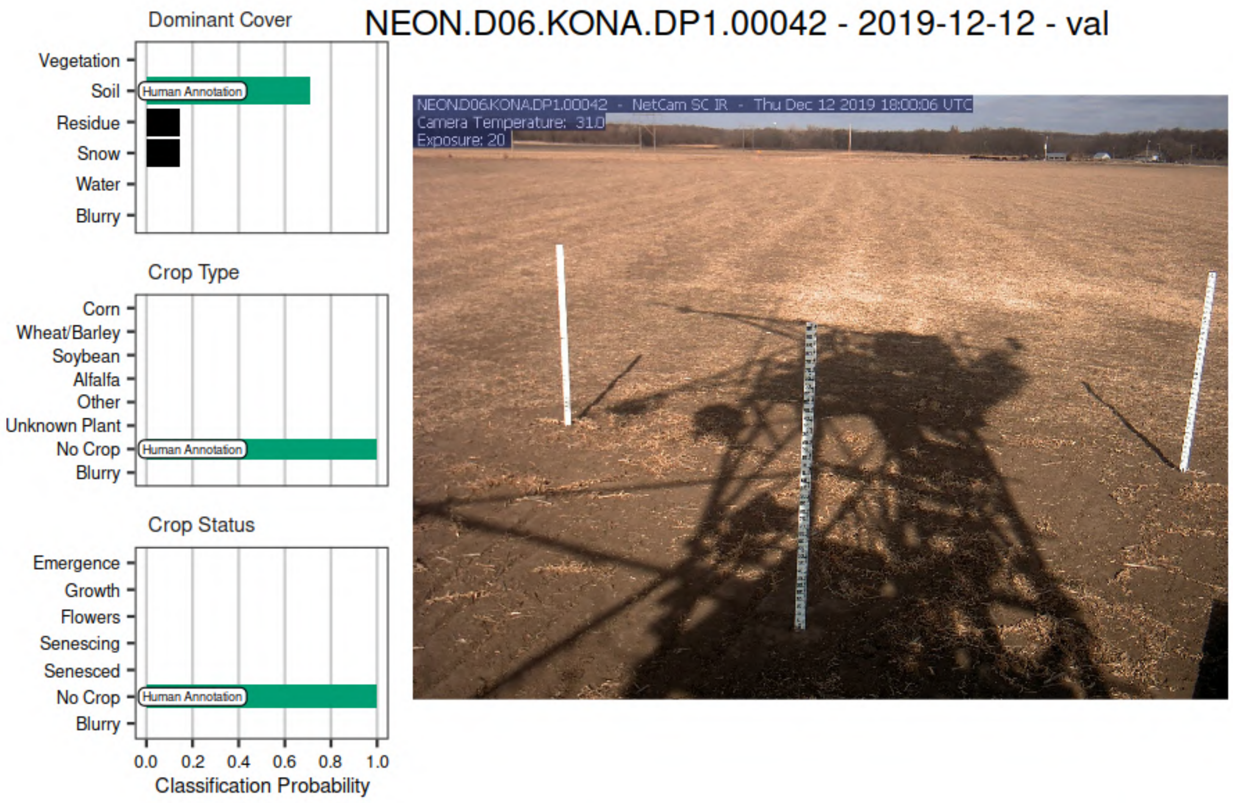


Figure S9

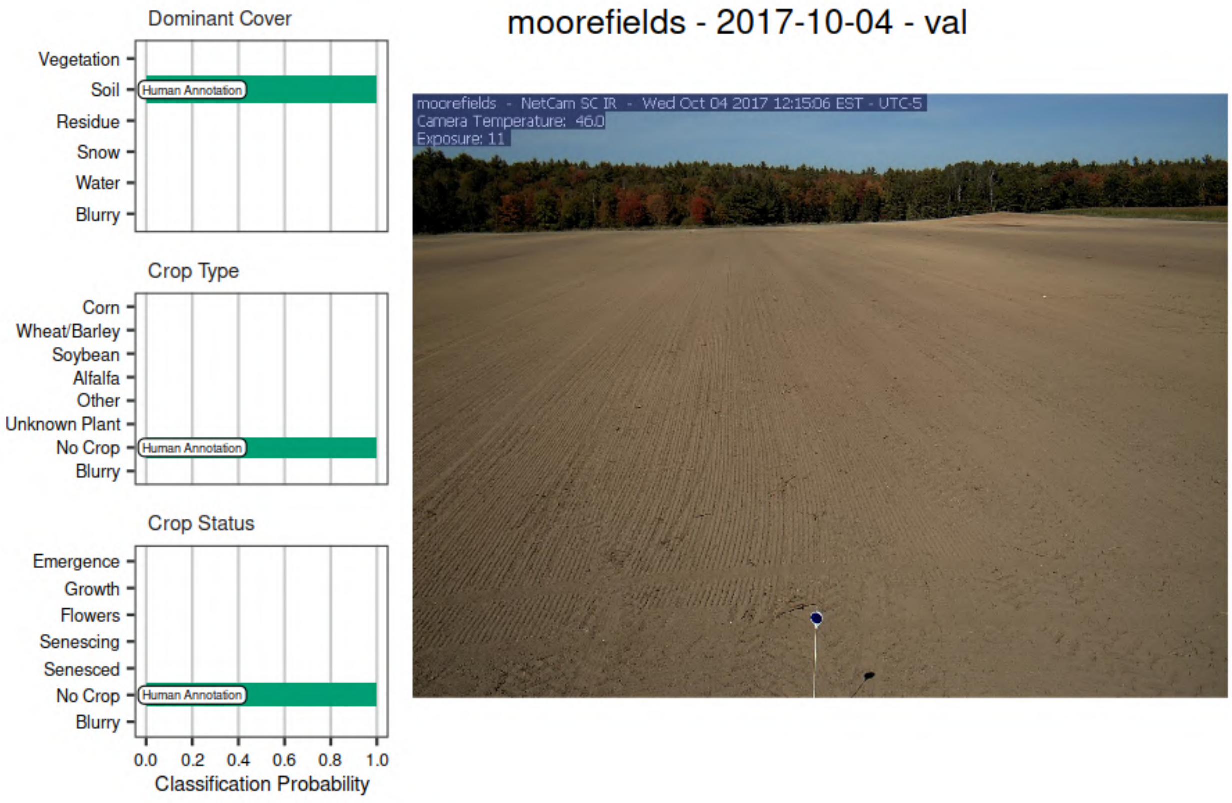


Figure S10

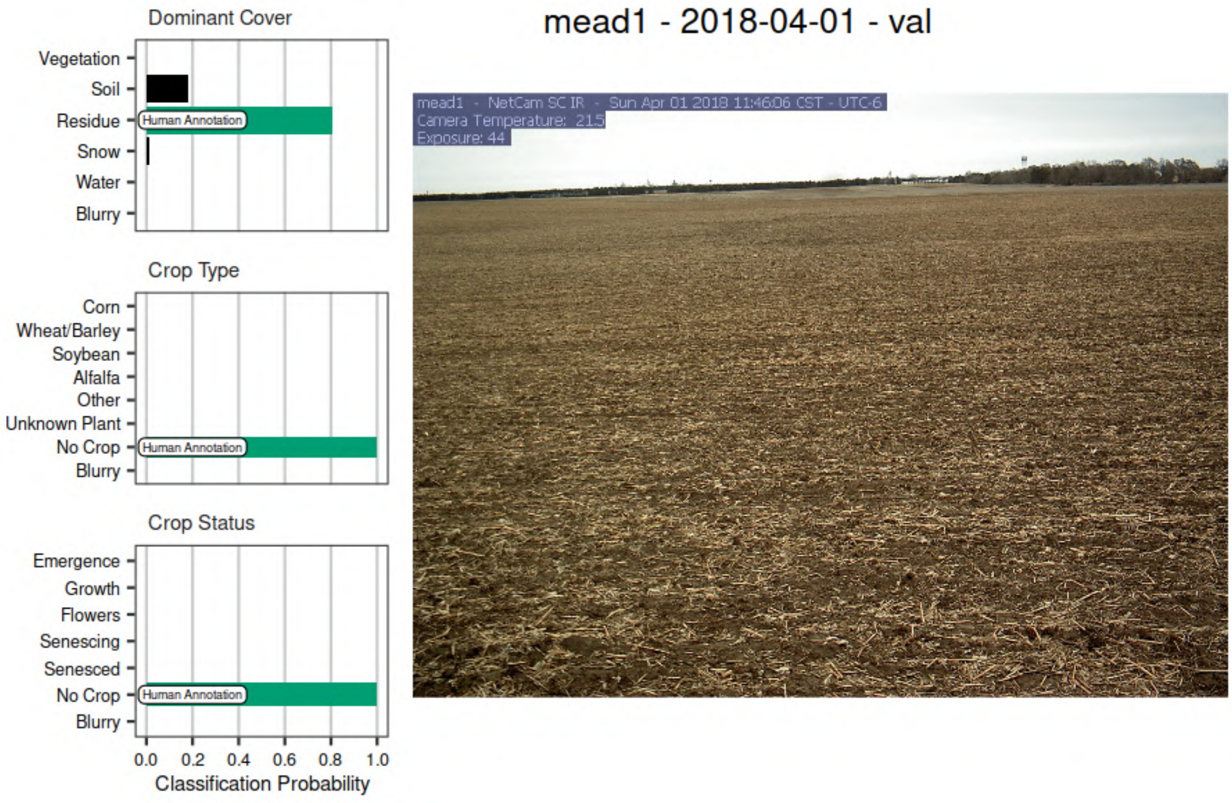


Figure S11

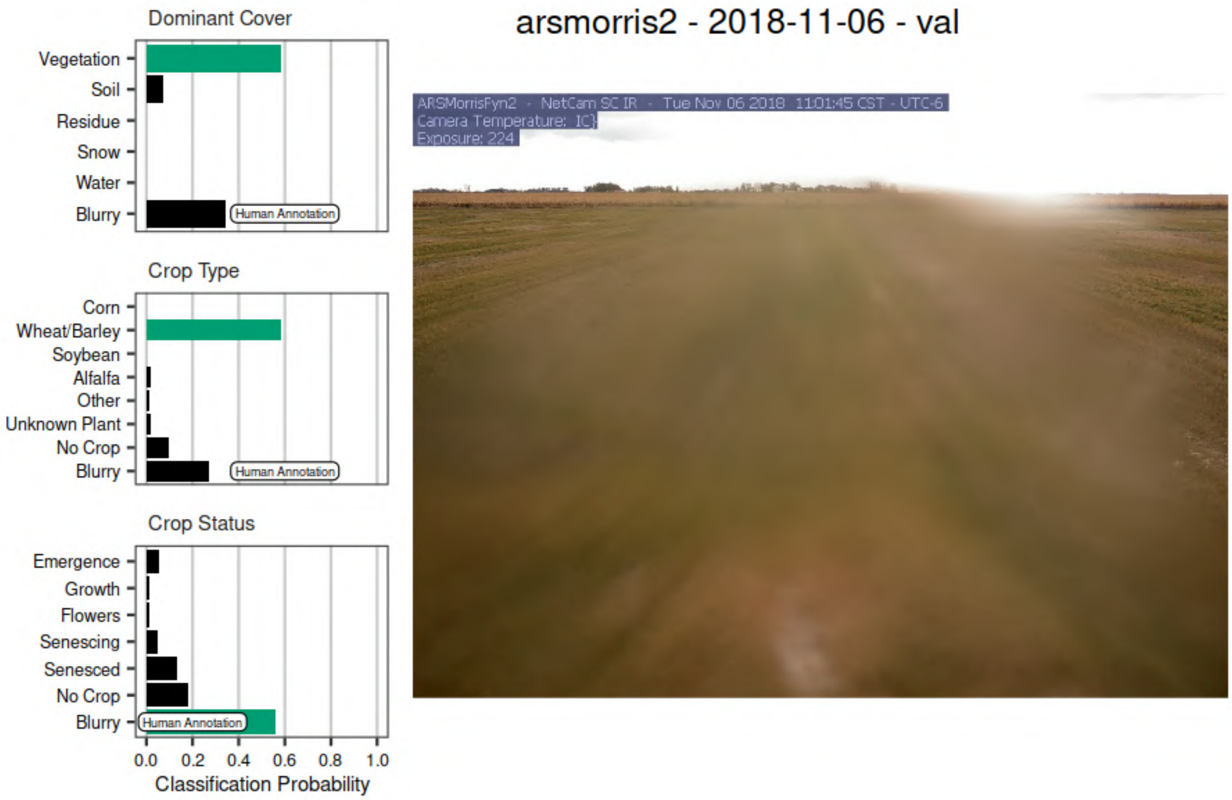


Figure S12

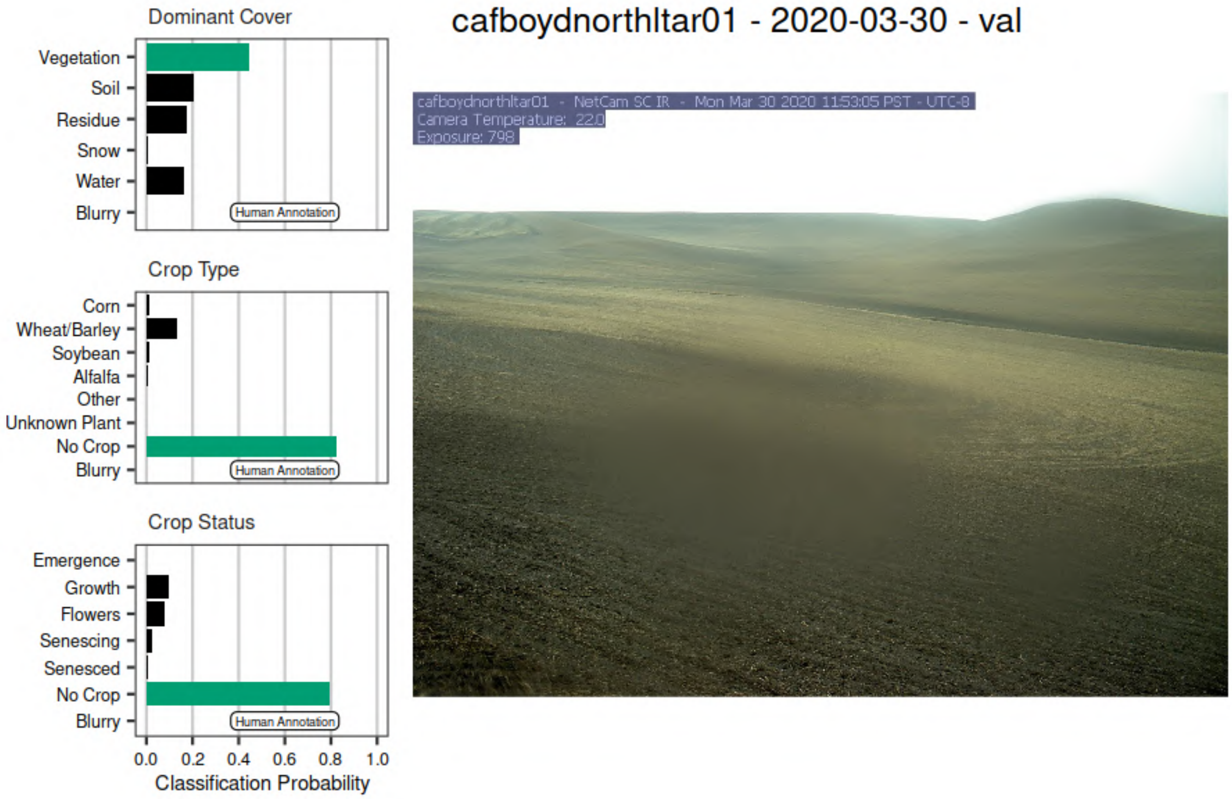


Figure S13

### Model Comparison

We originally used only the VGG16 model since it's initial performance was quite good, and comparable to similar studies. During the review process we performed a model comparison of several other CNN models pre-configured in the Keras library. The four models were VGG16, Xception, ResNet101V2, DenseNet169. We fit the models from scratch using all the same data described in the paper, except instead of 100k training images we used 15k so it finished in a reasonable time. Using a loss of categorical cross entropy we fit each model for 20 epochs using the Adam optimizer with learning rate of 0.01 and epsilon of 0.1. Below is a figure for validation loss over the 20 epochs for each model.

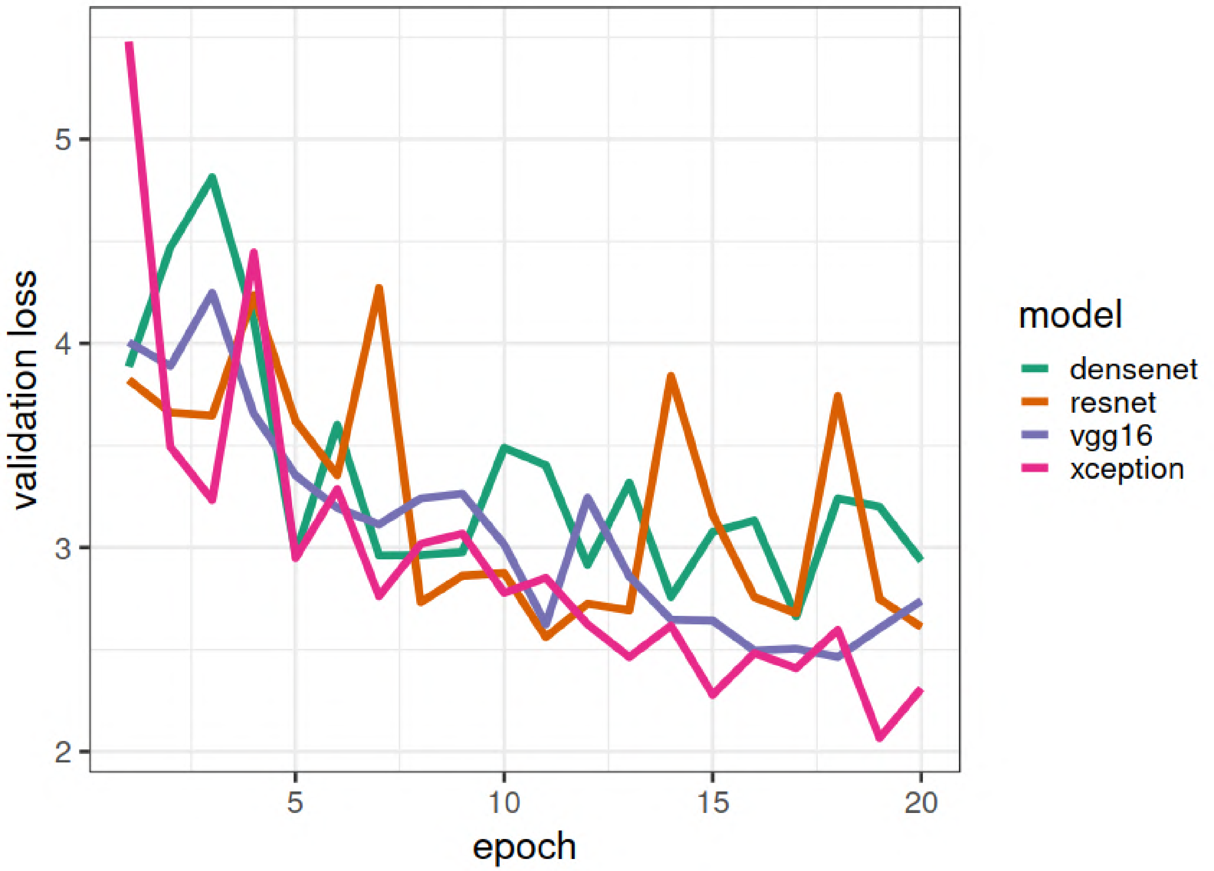


Figure S14: Trace of the validation loss for 20 epochs when comparing four different models using a subset of the training data.

The Xception and VGG16 models consistently performed the best. Thus, we did a final round of fitting the Xception model from scratch with the exact method described in the paper. We used the same 100k training images. We used the Adam optimizer and fit the Xception model for 15 epochs using a learning rate of 0.01 and epsilon 0.1, and an additional 5 epochs using a learning rate of 0.001 and epsilon of 0.1.  The following figure is the result for this fitting, in addition to the same fitting done for VGG16 in the original analysis.

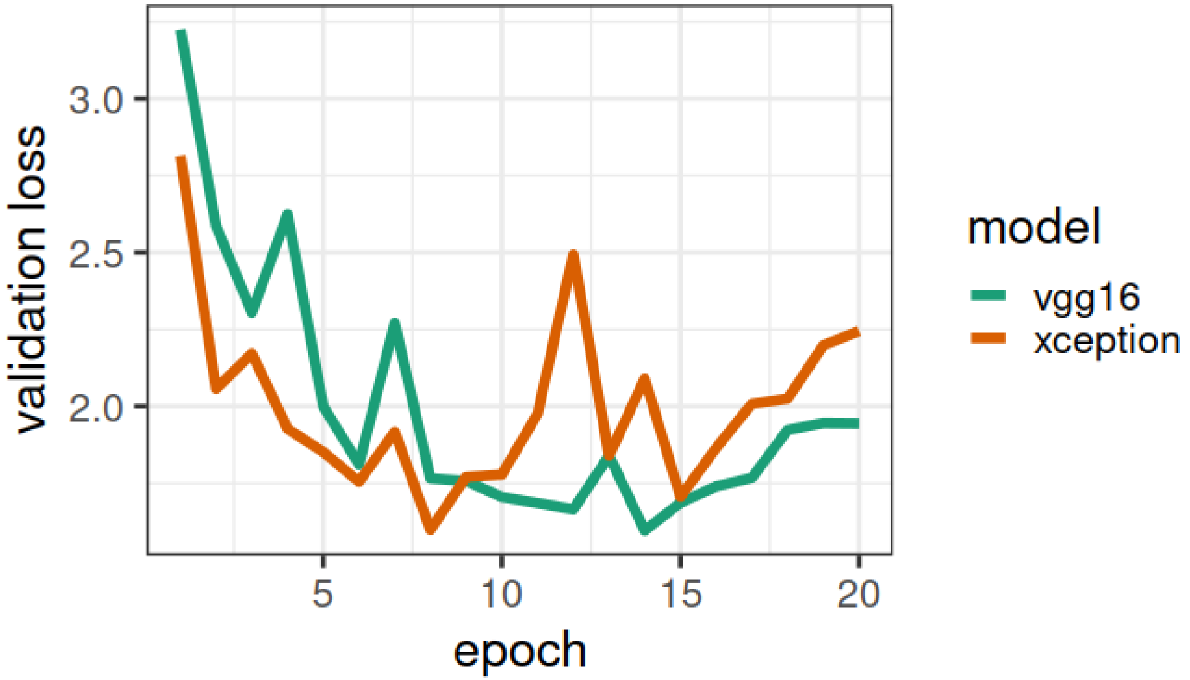


Figure S15: Trace of the validation loss for 20 epochs when comparing two of the best models using the full training set.

With the full training set the Xception model performed only as well as, or slightly worse, than the original VGG16 model. Therefore after the comparison we kept the VGG16 in place for the analysis.
